# Noradrenaline causes a spread of association in the hippocampal cognitive map

**DOI:** 10.1101/2024.08.31.610602

**Authors:** Renée S Koolschijn, Prakriti Parthasarathy, Michael Browning, Xenia Przygodda, Liliana P Capitão, William T Clarke, Tim P Vogels, Jill X O’Reilly, Helen C Barron

**Affiliations:** Wellcome Centre of Integrative Neuroimaging, University of Oxford, FMRIB, John Radcliffe Hospital, Oxford, United Kingdom; Radboud University, Donders Institute for Brain, Cognition and Behaviour, Nijmegen, the Netherlands; Department of Engineering, University of Cambridge, Cambridge, United Kingdom; University Department of Psychiatry, University of Oxford, Warneford Hospital, Oxford, United Kingdom; Oxford Health NHS Foundation Trust, Warneford Hospital, Oxford, United Kingdom; Medical Research Council Brain Network Dynamics Unit, Nuffield Department for Clinical Neurosciences, University of Oxford, Oxford, United Kingdom; German Center for Neurodegenerative Diseases (DZNE), Magdeburg, Germany; Institute for Cognitive Neurology and Dementia Research, Otto-von-Guericke University, Magdeburg, Germany; Psychological Neuroscience Lab, Psychology Researcher Centre (CIPsi), School of Psychology, University of Minho, Campus de Gualtar, Braga, Portugal; Institute of Science and Technology Austria, Klosterneuburg, Austria; Department of Experimental Psychology, University of Oxford, Oxford, United Kingdom

## Abstract

The mammalian brain stores relationships between items and events in the world as cognitive maps in the hippocampal system. However, the neural mechanisms that control cognitive map formation remain unclear. Here, using a double-blind study design with a pharmacological intervention in humans, we show that the neuromodulator noradrenaline elicits a significant ‘spread of association’ across the hippocampal cognitive map. This neural spread of association predicts overgeneralisation errors measured in behaviour, and is predicted by physiological markers for noradrenergic arousal, namely an increase in the pupil dilation response and reduced cortical GABAergic tone. Thus, noradrenaline sets the width of the ‘smoothing kernel’ for plasticity across the cognitive map. Elevated noradrenaline can allow disparate memories to become linked, introducing potential for distortions in the cognitive map that deviate from direct experience.

**Graphical abstract:** 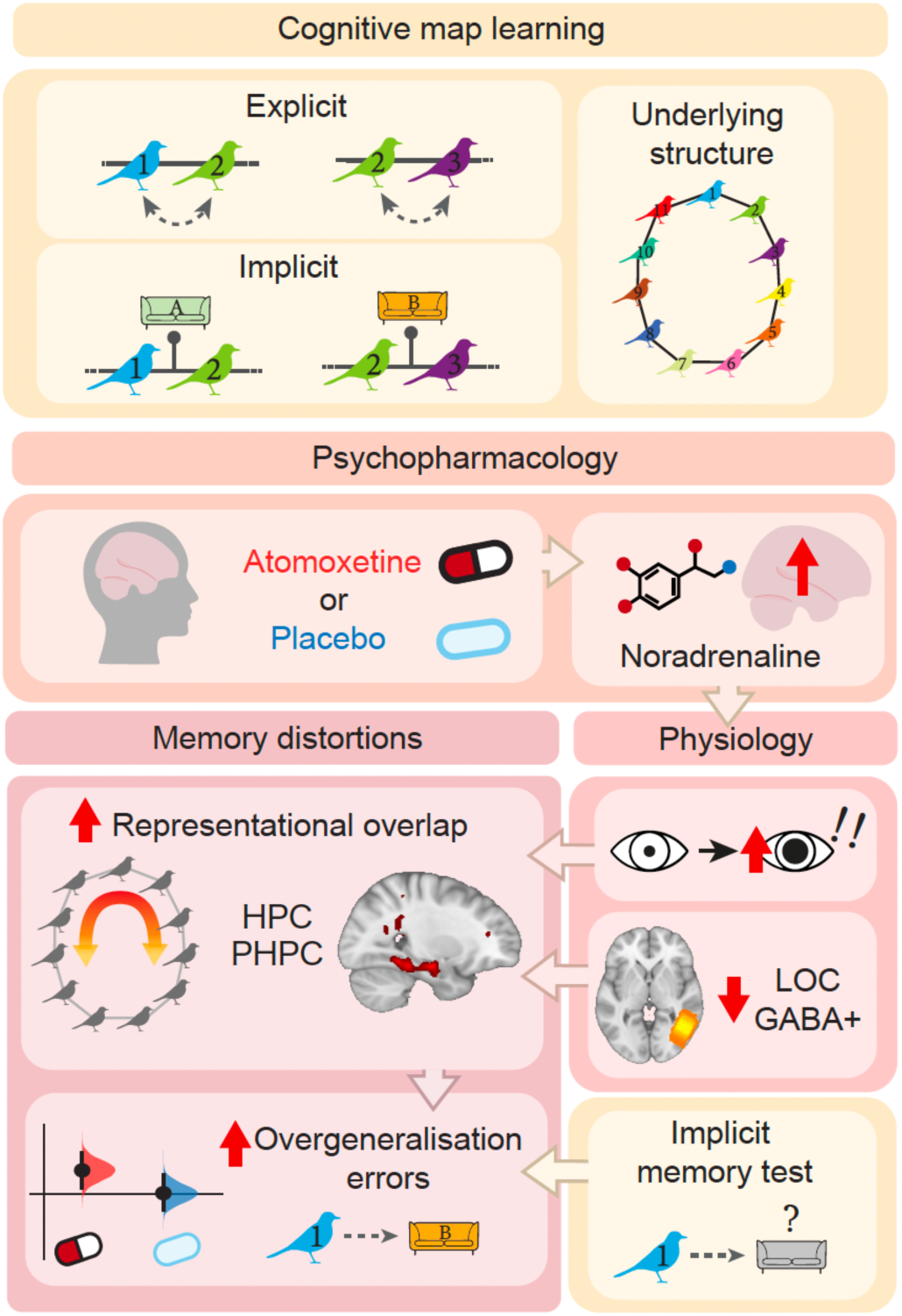

## Introduction

The mammalian brain stores the relationships between items and events that an organism experiences. This relational information is represented by the hippocampal system, as a cognitive map of the environment (O’Keefe & Nadel, 1978; Tolman, 1948). Following its formation, a cognitive map can be used to recall information from the past, while also providing a basis to predict the consequences of upcoming actions and guide future behaviour (Behrens et al., 2018; Epstein et al., 2017). Critically, a cognitive map can extend beyond direct experience, representing more than the sum of its parts by including inferred relationships between stimuli or events in the world (Barron et al., 2020; Gupta et al., 2010; Schwartenbeck et al., 2023; Wu & Foster, 2014). These inferred relationships can be used to draw associations that we have not directly experienced, to support inferential choice (Barron et al., 2020). For this reason, cognitive maps are credited with enabling adaptive and flexible behaviours (Behrens et al., 2018).

Yet, there is a fine balance to be struck when building a cognitive map that extends beyond direct experience. On the one hand, a cognitive map can support our ability to take new shortcuts and infer unobserved relationships between different events (Barron et al., 2020; Dragoi & Tonegawa, 2011; Ólafsdóttir et al., 2015). On the other hand, by deviating from reality a cognitive map has the propensity to introduce unwanted distortions in memory (Bellmund et al., 2020; McNamara, 1986; Warren et al., 2017). Thus, the neural mechanisms responsible for forming a cognitive map critically determine whether the cognitive map facilitates adaptive or maladaptive behaviour. However, the neural mechanisms which control cognitive map formation remain unknown, including the precise mechanism that titrates the extent to which discrete stimuli or events become associated (the ‘spread of association’) within a cognitive map.

One candidate mechanism for titrating the ‘spread of association’ within the cognitive map involves neuromodulators, including the neuromodulator noradrenaline (Atherton et al., 2015; Mather et al., 2016; Sara, 2009). Noradrenaline profoundly alters plasticity in the brain by reducing the induction threshold for synaptic plasticity and reducing activity in inhibitory interneurons (Hagena et al., 2016; Lemon et al., 2009; Lisman & Grace, 2005; Martins & Froemke, 2015; Stringer et al., 2016; Tsetsenis et al., 2022). As a result, noradrenaline shifts the balance between cortical excitation and inhibition (E/I) in favour of excitation (Froemke, 2015). Learning during periods of elevated neuromodulatory tone therefore facilitates excitatory plasticity, not only strengthening individual memories but also providing an opportunity for ‘runaway excitation’ to link discrete memories in a cognitive map (Cai et al., 2016; Schapiro et al., 2013). In the extreme, associations may form between memories that have not been experienced together but reside in close proximity in the cognitive map. In this manner, noradrenaline may render new memories prone to overgeneralisation or distortion.

To test the hypothesis that noradrenaline titrates ‘spread of association’ between memories within the cognitive map, we used a combination of computational and experimental approaches. First, we embedded memories into a spiking neural network model with biologically plausible plasticity at excitatory and inhibitory synapses. This model predicted a spread of association in the underlying memory map when learning occurs during a period of reduced inhibitory tone, as expected during elevated noradrenaline. We then tested these predictions empirically, using a double-blind study design with a pharmacological intervention in humans. We show that the neuromodulator noradrenaline elicits a significant ‘spread of association’ across the cognitive map in hippocampus and parahippocampus. Crucially, the neural spread of association is predicted by physiological markers for noradrenergic arousal, namely an increase in the pupil dilation response and a reduction in cortical inhibition. Further, we show that the neural spread of association predicts overgeneralisation errors in memory measured in behaviour 4 days after learning. Together these findings demonstrate that elevated noradrenaline allows disparate memories to become linked, introducing potential for distortions in the cognitive map that deviate from direct experience. Therefore, by setting the width of the ‘smoothing kernel’ for plasticity, levels of noradrenaline during learning control the extent to which memories become associated within a cognitive map.

## Results

### Experimental design

We predicted that learning during elevated noradrenaline should increase the spread of association in the underlying cognitive map. To test this prediction, we manipulated noradrenaline during a learning task using a between-subject, double-blind, randomised placebo-controlled design. At the start of the experiment, participants were given either a single dose of the noradrenergic reuptake inhibitor, atomoxetine (‘ATX’), or a placebo (‘PLC’). Previous studies demonstrate that atomoxetine can be used to increase levels of noradrenaline throughout the brain (Bymaster et al., 2002; Easton et al., 2007). Our blinding procedure was effective and participants in the drug group (ATX) were unable to correctly report whether they received atomoxetine or placebo (Supplementary Figure 1, Supplementary Table 1).

After receiving atomoxetine or placebo, participants waited 90 minutes to allow noradrenaline levels to reach peak plasma concentration (Sauer et al., 2005). Participants then completed a learning task. We designed the task such that the underlying task structure was susceptible to overgeneralisation and memory distortion. The task required participants to learn a set of pairwise associations that could be arranged in a ring structure (Figure 1A). Participants were not shown the ring structure but were instead trained on 11 pairwise associations that made up the ring. Each *pairwise association* was a pair of bird stimuli, with each pair presented in a specific context indicated by a living room interior (Figure 1B-C, Supplementary Figure 2). In addition, each bird stimulus was paired with two other bird stimuli such that each *bird* appeared in two different contexts, depending on its pairing. For example, bird 2 appeared both with bird 1 (in room A) and with bird 3 (in room B). Participants were not made explicitly aware of the underlying task structure. Memory errors between bird stimuli and contextual cues provided a means to later test participants’ susceptibility to overgeneralise and distort the relational task structure.

**Figure 1.**
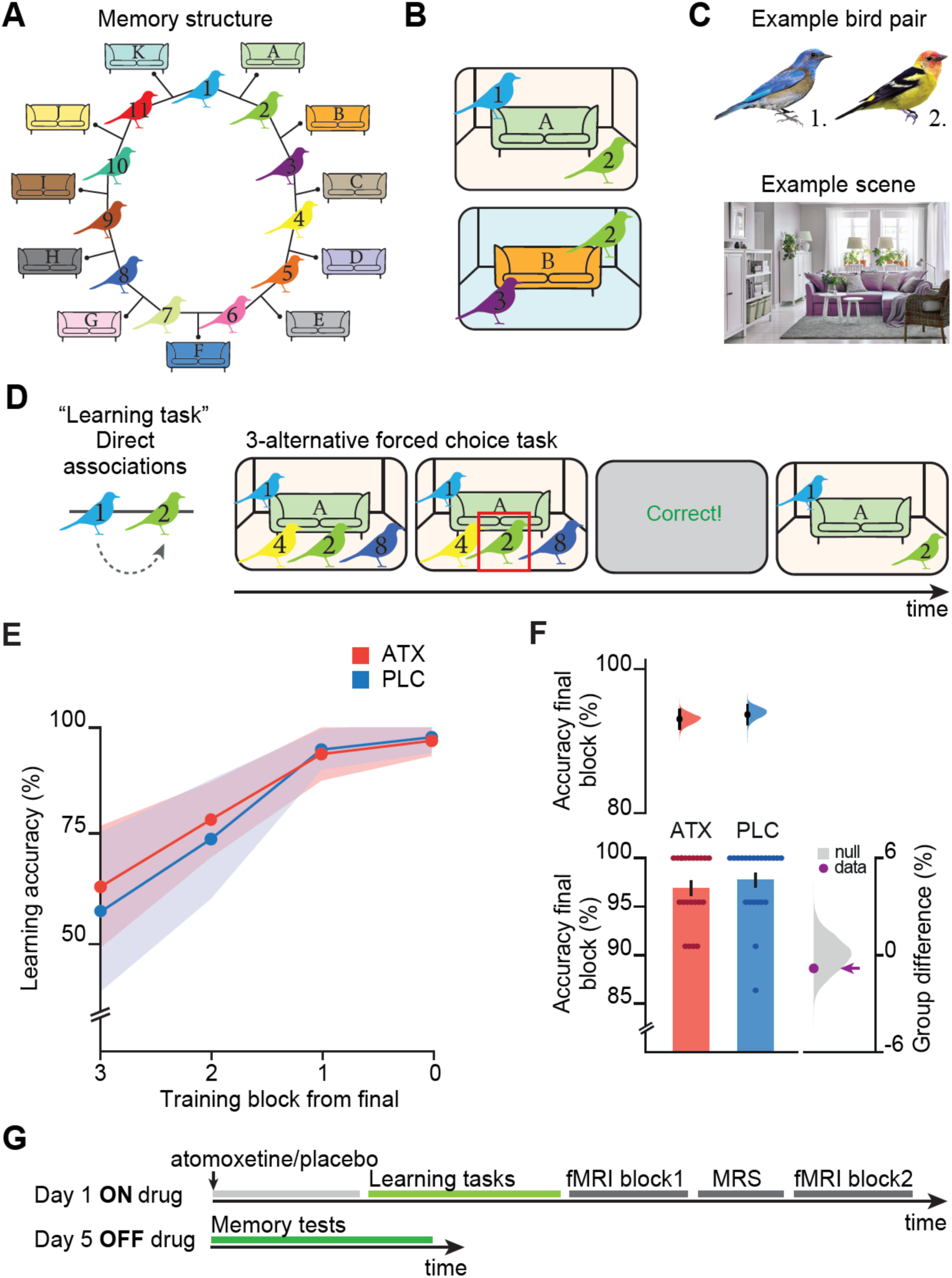
Experimental design used to assess the effect of ATX on forming a cognitive map. **A-B** Schematic: participants explicitly learned associations between pairs of birds. Each pair of birds was shown against a particular contextual cue, with implicit exposure. Overall, each bird stimulus was associated with two other bird stimuli, such that all stimuli could be arranged as a ring structure, although this information was not explicitly shared with participants. **C** Example pair of bird stimuli together with an example contextual cue which includes a living room scene containing a sofa. **D** To learn the context-dependent pairwise associations, on day 1 participants completed a three-alternative forced choice task with feedback (‘learning task’). Participants were tested on the bird-bird associations, as indicated by the grey dotted arrow**. E** Participants performed the learning task until they successfully remembered >90% of the associations. **F** All participants successfully completed the learning task, with no significant difference between groups (ATX – PLC: permutation test: p=0.206). Upper right: Bootstrap-coupled estimation (DABEST) plots. Black dot, mean; black ticks, 95% confidence interval; filled curve, sampling error distribution. Lower right: memory accuracy (mean +/− SEM). Lower right: null distribution of the group differences generated by permuting subject labels, purple dot: true group difference. **G** Schematic of the experimental protocol. On Day 1, participants received either atomoxetine or placebo. After a 90 min waiting period participants completed the learning tasks and underwent an MRI scan. On day 5, participants completed memory tests.

Participants learned to associate pairs of birds using a three-alternative forced choice task (‘Learning task’, Figure 1D). During the learning task participants were not tested on the relationships between the bird stimuli and the contextual cues, leaving these relationships implicit. All participants learned the pairwise associations with high accuracy and there was no significant difference in learning between the ATX and PLC groups (Figure 1E-F, Supplementary Figure 3B). After learning, participants were given the opportunity to infer indirect relationships between bird stimuli that share a common association, thereby making trajectories around the ring structure (‘inference task’, without feedback; Supplementary Figure 3A).

Importantly, this task design allowed us to test whether elevated noradrenaline introduces a spread of association across the underlying cognitive map. Specifically, we predicted that if noradrenaline increases runaway excitation in the underlying cognitive map for the ring of associations (Figure 1A), stimuli and contextual cues situated in proximal locations on the ring may become erroneously associated due to a ‘spread of association’. By contrast, those situated at distal locations on the ring should remain distinct, thus providing the necessary control for those in proximal locations. Our experimental design thus provided a precise and controlled means to test evidence for the effect of noradrenaline on forming a cognitive map, at both a behavioural and neural level.

### Behavioural evidence for overgeneralisation errors with noradrenaline

Four days after learning (Day 5, Figure 1G), when the effects of atomoxetine had washed out (Sauer et al., 2005), participants returned to perform memory tests designed to provide behavioural evidence for overgeneralisation across memories. We included memory tests for both the pairwise associations between birds learned on day 1 (‘explicit memory test’, Figure 2A) and the incidental pairings between the bird stimuli and contextual cues (‘implicit memory test’, Figure 2B).

**Figure 2.**
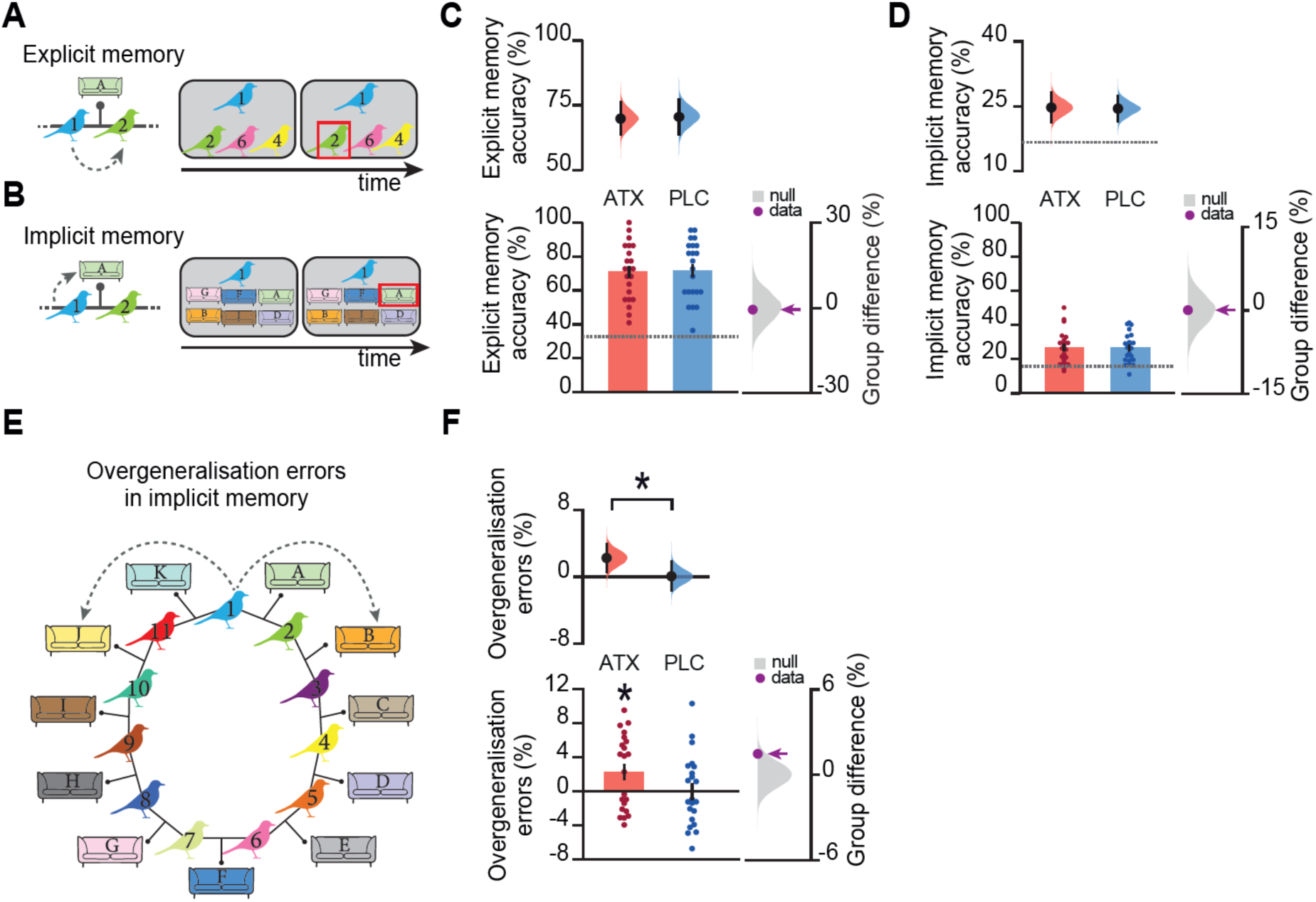
Elevated noradrenaline during learning increases overgeneralisation in behaviour. **A** Example trial of the explicit memory test performed by participants on day 5, where the participants were tested on the bird-bird associations learned on day 1, as indicated by the grey dotted arrow. **B** Example trial of the implicit memory test performed by participants on day 5, where the participants were required to recall which contextual cue (sofa) was incidentally paired with each bird stimulus, as indicated by the grey dotted arrow. **C**-**D,F** Upper: Bootstrap-coupled estimation (DABEST) plots. Black dot, mean; black ticks, 95% confidence interval; filled curve, sampling error distribution. Lower left: memory accuracy (mean +/− SEM). Lower right: null distribution of the group differences generated by permuting subject labels, purple dot: true group difference. **C** In the explicit memory test, overall memory accuracy was high and there was no significant difference in overall accuracy between groups (ATX – PLC: permutation test: p=0.421). **D** In the implicit memory test, overall accuracy was low. There was no significant difference in overall accuracy between groups (ATX – PLC: permutation test: p=0.498), but the relatively high proportion of error trials afforded an opportunity to test evidence for overgeneralisation, as shown in *E-F*. **E** Schematic illustrating the predicted distortion in the underlying cognitive map, where a ‘spread of association’ leads to overgeneralisation in participants’ implicit memory such that errors in recall are biased towards contextual cues that are paired with incorrect but proximal bird stimuli on the underlying ring structure, as indicated by the grey dotted arrow. **F** During the implicit memory test shown in *B*, participants made errors on 75.4% of the trials (ATX: 75.3%, PLC: 75.6%). ‘Overgeneralisation errors’ were defined as those errors where the incorrect chosen contextual stimulus was proximal to the probe, as shown in *E*. The ATX group made a significant number of overgeneralisation errors (p=0.014, effect size computed from 10,000 bias-corrected bootstrapped resamples (Ho et al., 2019)), and significantly more overgeneralisation errors compared to PLC (ATX – PLC: permutation test p=0.048).

For behavioural tests on day 5 where overall accuracy in both groups was relatively high, we observed no significant group difference in memory accuracy (ATX vs. PLC, ‘explicit memory accuracy’ and ‘inference accuracy’, Figure 2C, Supplementary Figure 3C). However, for the implicit memory test, where overall accuracy in both groups was relatively poor, the error trials (74% of trials, on average) afforded an opportunity to test evidence for overgeneralisation and distortions in memory.

Specifically, we reasoned that if the underlying memory map is distorted to include a ‘spread of association’ around the ring, participants should be more likely to overgeneralise by associating bird stimuli with incorrect contextual cues that are proximal rather than distal on the ring. For example, for bird 1, rather than selecting room A where bird 1 and bird 2 were located, participants made an overgeneralisation error if they selected room B, where bird 2 and bird 3 were located. We hypothesised that learning under noradrenaline should increase these overgeneralisation errors, due to the spread of association in the underlying cognitive map. Consistent with this prediction, the ATX group compared to PLC was significantly more likely to erroneously report associations between proximal rather than distal stimuli on the ring (Figure 2F). These results suggest that learning under elevated noradrenaline increases the spread of associations between stimuli that were never presented together but are located nearby in a cognitive map.

### Physiological markers of elevated noradrenaline

In clinical trials substantial interindividual variability is observed in response to neuropsychiatric medication, such as atomoxetine. Here, we measured this variability to quantify interindividual differences in the response to drug in the ATX group. We used two physiological markers of noradrenergic arousal, namely the pupil dilation response, and cortical measures of GABAergic tone.

#### Pupil response to oddball stimuli is elevated in ATX

Work in animals and humans shows that increases in pupil diameter reflect increases in the availability of noradrenaline (Gilzenrat et al., 2010; Joshi et al., 2016; Rajkowski et al., 1993; Reimer et al., 2016; Samuels & Szabadi, 2008). Here we took advantage of this physiological marker for noradrenaline. We collected pupillometry data after the learning task during an “oddball” detection task performed inside the MRI scanner. On each trial participants were presented with either a familiar stimulus (91% of trials) previously observed during the learning task, or an oddball stimulus (9% of trials, Figure 3A). Participants were required to make a button press response when they detected an oddball stimulus. We analysed pupil dilation following stimulus presentation. Consistent with previous studies (Nassar et al., 2012; O’Reilly et al., 2013; Preuschoff et al., 2011) and in line with the suggestion that noradrenaline signals ‘unexpected uncertainty’ (Yu & Dayan, 2005), in both the ATX and PLC groups there was an enlarged phasic pupil response following oddball stimuli. This oddball response was greater in the ATX group compared to the PLC group, with a significant group difference from ~6-10 s after stimulus onset (Figure 3B). Thus, the ATX group showed an increase in their sustained pupil dilation response to surprising stimuli, providing a physiological maker for the drug response of individuals within this group.

**Figure 3.**
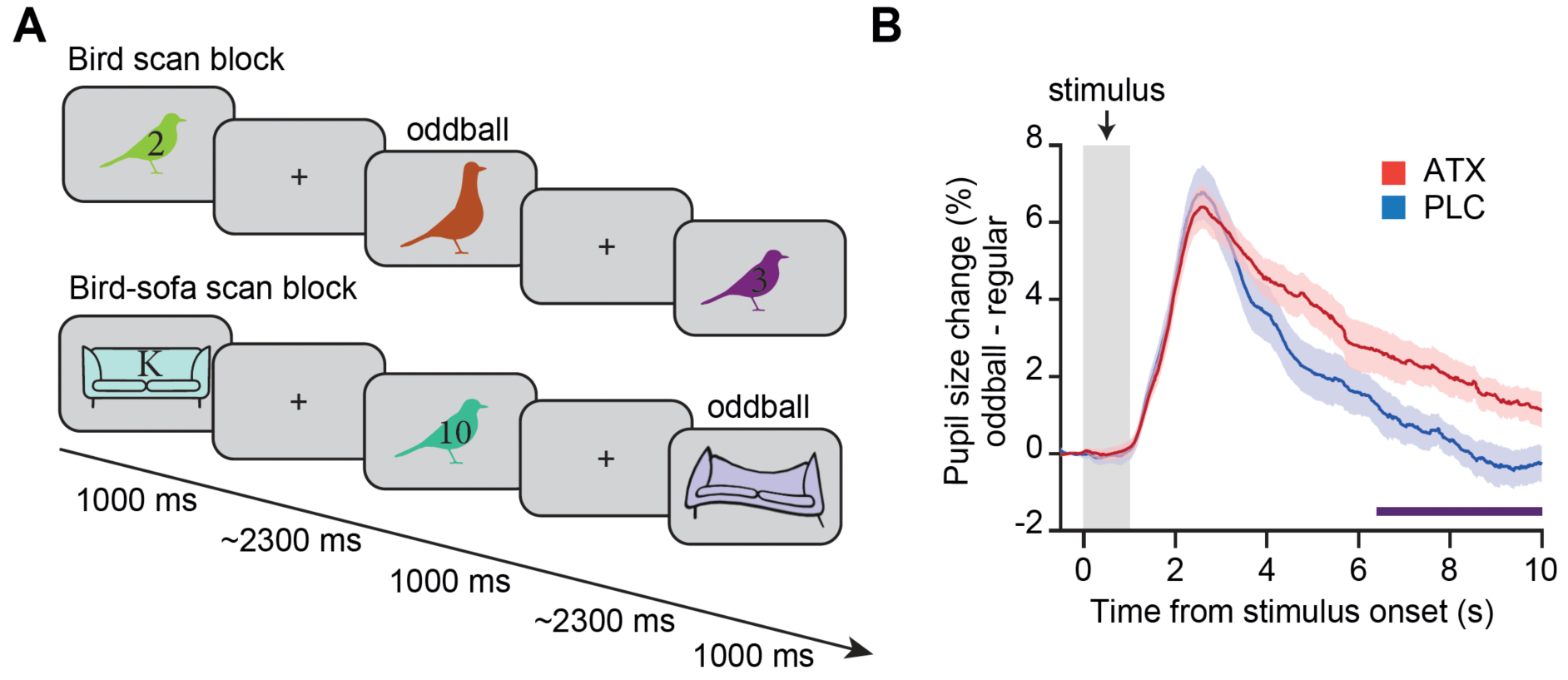
Elevated noradrenaline increases pupil surprise response. **A** During the scan task, participants observed bird and sofa stimuli encountered during the learning task presented against a grey background. Participants were instructed to detect “oddball” stimuli, encountered in ~9% of trials, which constituted warped or differently coloured versions of the familiar stimuli. **B** Pupil dilation was larger in response to oddball compared to familiar stimuli, in both the ATX and PLC groups. Between 6 and 10 s after stimulus onset, the ATX group showed a significantly larger pupil dilation response to oddball stimuli (purple horizontal line: cluster-corrected significant ATX – PLC; permutation test corrected for multiple comparisons: p=0.025).

#### Elevated noradrenaline decreases inhibitory tone in neocortex

Having established that a single dose of atomoxetine increases pupil response to surprising stimuli, we tested evidence for a second physiological marker of elevated noradrenaline, namely cortical inhibition. The majority of noradrenergic projections to cortex originate in the locus coeruleus (LC) and stimulation of LC in rodents induces a decrease in cortical inhibitory tone (Martins & Froemke, 2015). In humans, a single dose of atomoxetine reduces paired-pulse suppression using visual evoked potentials in visual cortex, indicating an increase in cortical excitability (Höffken et al., 2012). Taken together, we predicted that elevating noradrenaline using a single dose of atomoxetine should induce a reduction in cortical inhibition, to shift cortical activity in the direction of increased excitability.

To measure cortical inhibition in our participants, we used Magnetic Resonance Spectroscopy (MRS) (Figure 1G) to quantify the local concentration of different neural metabolites in the brain, including the concentration of cortical gamma-aminobutyric acid (GABA+, GABA and macromolecules) together with Glx (glutamate and glutamine). In line with previous evidence from rodents (Martins & Froemke, 2015), GABA+ measured from a voxel positioned in the LOC (Figure 4A), a visual object processing region (Grill-Spector et al., 1999, 2001), was significantly reduced in the ATX group compared to the PLC group (Figure 4C). Moreover, this reduction in GABA+ in the ATX group accounted for a significant increase in the ratio between Glx and GABA+ in the ATX compared to the PLC group (Glx:GABA+, Figure 4D) (Koolschijn et al., 2021). Critically, there was no significant difference in MRS quality metrics between the ATX and PLC groups (Supplementary Figure 5). When measuring GABA+ and Glx:GABA+ ratio from V1, no significant difference between ATX and PLC was observed (Supplementary Figure 6). The lack of effect in V1 may be explained by lower Signal to Noise Ratio (SNR) in V1 compared to LOC, as indicated by significantly lower temporal SNR (tSNR) in this region (Supplementary Figure 7). In LOC, significant evidence for reduced GABA+ in ATX compared to PLC suggests that a single dose of atomoxetine reduces cortical inhibition in brain regions sensitive to task stimuli, namely the visual object processing region LOC. Moreover, taking Glx:GABA+ as a proxy measure of the balance between cortical excitation and inhibition (E/I), these results suggest that noradrenaline shifts participants’ E/I ratio along a continuum in the direction of increased cortical excitability. Taken with the pupil dilation response, this MRS measure of reduced cortical inhibition or E/I ratio in ATX provides a second physiological marker of the response to drug.

**Figure 4.**
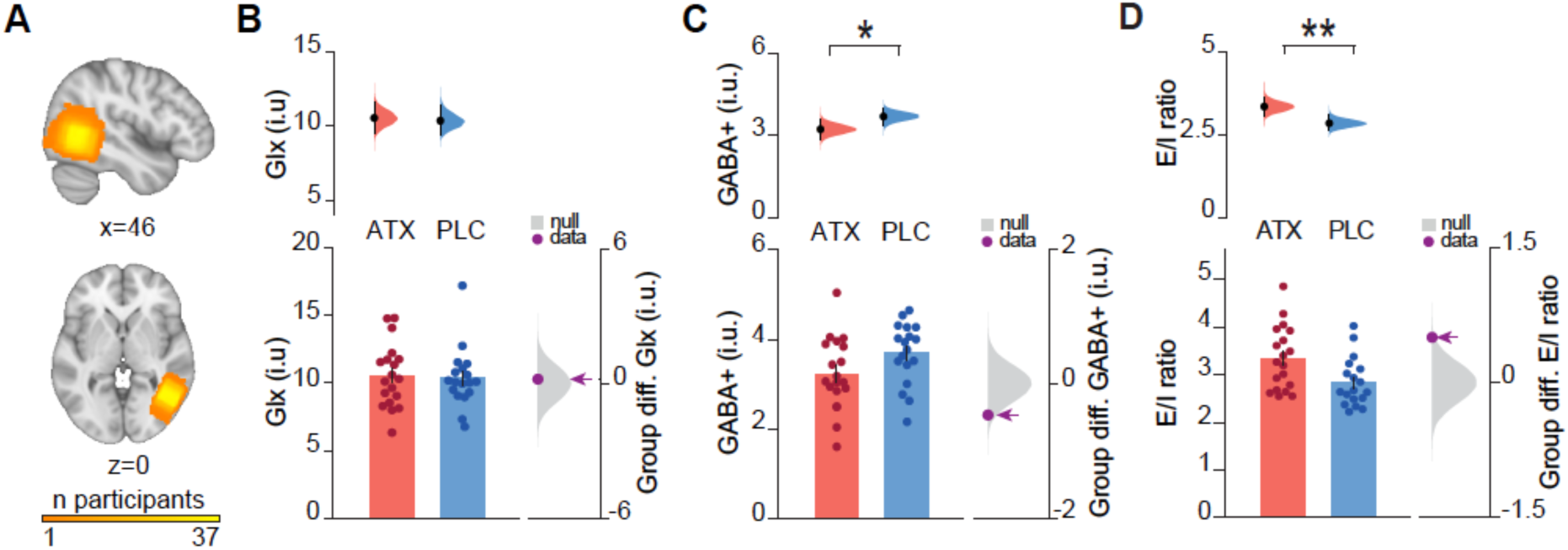
Elevating noradrenaline decreases inhibitory tone in LOC. **A** Anatomical location of the 2 x 2 x 2 cm^3^ MRS voxel, positioned in LOC. Cumulative map across 37 participants. **B-D** We used MRS to quantify the effect of atomoxetine on excitatory and inhibitory tone in neocortex, ~3.5 h after drug intake. Upper: Bootstrap-coupled estimation (DABEST) plots. Black dot, mean; black ticks, 95% confidence interval; filled curve, sampling error distribution. Lower left: Glx (***B***) and GABA (***C***) concentrations, and E/I ratio (***D***) (mean +/− SEM). Lower right: grey: null distribution of the group differences; purple dot: true group difference. **B** No significant group difference was observed between ATX and PLC for the concentration of Glx (ATX – PLC: permutation test p=0.402). **C** The concentration of GABA+ was significantly lower in ATX compared to PLC (ATX – PLC: permutation test p=0.034). **D** E/I ratio, defined as the ratio of Glx:GABA+, was significantly greater in ATX compared to PLC (ATX – PLC: permutation test p=0.007).

### Neural network model shows that reduced inhibitory plasticity during learning leads to a spread of association effect

Having confirmed that a single dose of atomoxetine induces expected changes in physiological markers of noradrenergic arousal, we next investigated the effect of elevated noradrenaline on the formation of a cognitive map of the task. At a behavioural level, overgeneralisation errors were observed in ATX but not PLC (Figure 2F). Here, we asked whether this behavioural evidence for overgeneralisation in the ATX group can be accounted for by distortions in the underlying neural representation of the task. Specifically, we predicted that learning under elevated noradrenaline should introduce a systematic spread of association in the underlying memory map, due to local ‘runaway excitation’ around the ring structure, where proximal nodes show more representational overlap compared to distal nodes. To test this hypothesis we used a combination of computational and experimental approaches.

First, we built a computational model to probe whether learning during a period of reduced inhibition alters synaptic modification to allow for a spread of association in the underlying cognitive map. To test this hypothesis we refined a set of previously published spiking neural network models (Agnes & Vogels, 2024; Barron et al., 2016; Vogels et al., 2013) and replicated our experiment using a reduced ring of associations in silico. To represent distinct stimuli we first embedded six independent and nonoverlapping cell assemblies, or ‘nodes’ *A-F,* that were balanced by local inhibition. To simulate the memory map in its pre-learning state (prior to learning the ring of associations), we externally activated one independent cell assembly ‘*A*’ (to simulate sensory input) and found no change in the activity of other nonoverlapping assemblies (Figure 5A).

**Figure 5.**
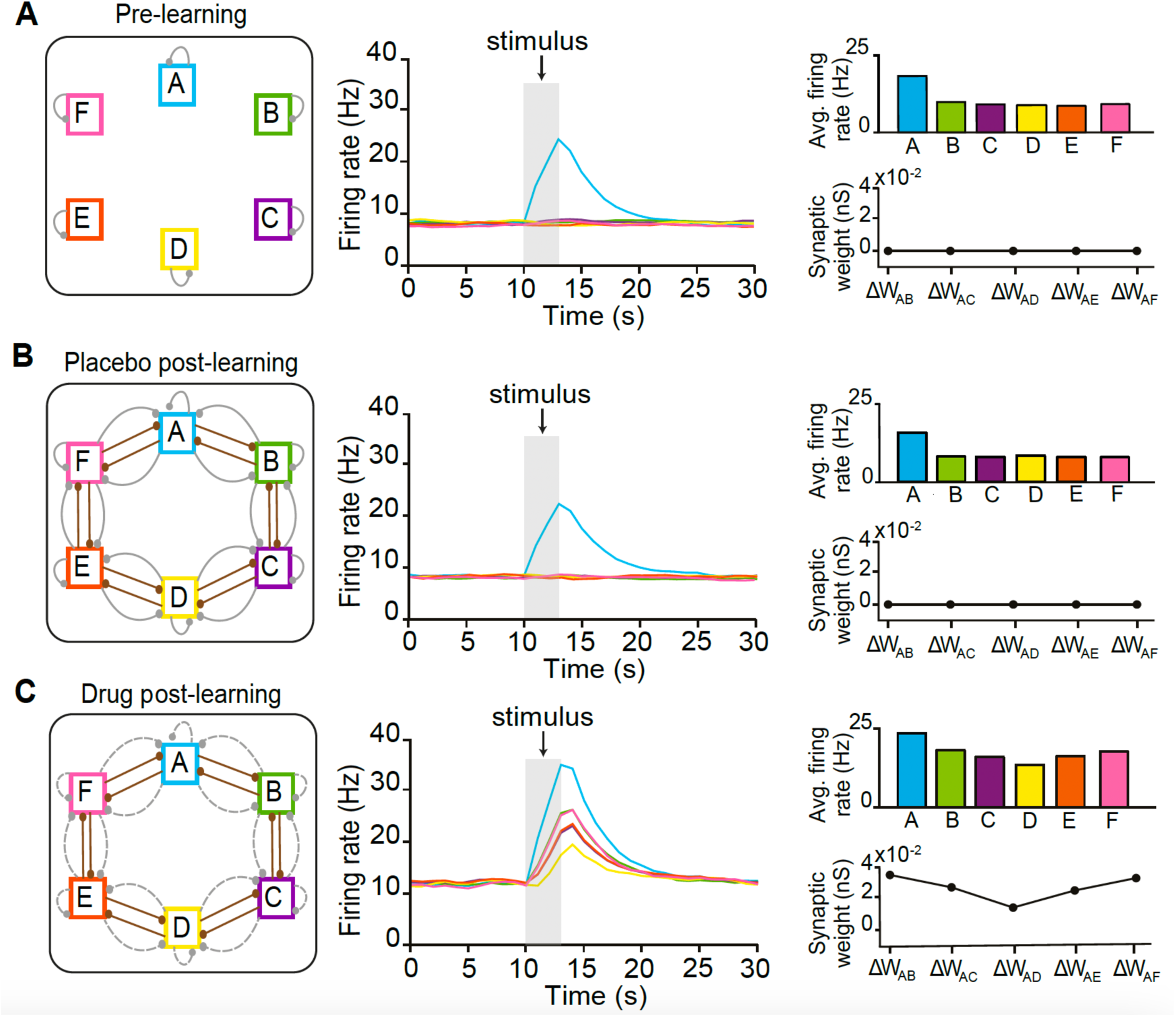
Neural network model showing how elevated noradrenaline during learning can lead to a spread of association in the underlying memory map. **A-B** Three snapshots of the recurrent spiking neural network, each showing the response of the network to stimulating embedded cell assembly ‘*A*’. Left: Schematic showing the architecture and parameter conditions of the network. Six cell assemblies are pictured as coloured squares. Excitatory and inhibitory connections are drawn in brown and grey, respectively. Dotted lines indicate weaker connections. Middle: Average firing rate of all excitatory neurons in each assembly, in response to activation of assembly ‘*A*’ via externally driven input during the period marked “stimulus”. Right Upper: Average firing rates of the excitatory neurons in each assembly during the “stimulus” period. Right Lower: Change in the mean synaptic weight between assembly ‘*A*’ and other assemblies following activation of assembly ‘*A*’. **A** In the pre-learning state, activation of cell assembly ‘*A*’ leads to high firing rates in the activated neuron group, but not in the other assemblies. **B-C** Post-learning state, after the ring of associations had been embedded by strengthening the excitatory (E-E) connections between neighbouring cell assemblies. **B** Under ‘placebo’, activation of cell assembly ‘*A*’ still selectively led to high firing rates in cell assembly ‘*A*’ only. Activity in all the other assemblies remains uniform, with no change in the conductances of synapses between ‘*A’* and any other assembly. **C** Under ‘atomoxetine’, activation of cell assembly ‘*A*’ led to high firing rates in the activated neuron group, but also resulted in graded co-activation across neurons in the other assemblies. The graded co-activation in other assemblies occurred relative to their respective distances from assembly *‘A*’. A gradient can also be observed across the average firing rates of each assembly during the “stimulus” period (upper right). The reduced firing in inhibitory neurons promotes excitatory plasticity, which also reflects a gradient with greater change in synaptic conductances between assembly ‘*A*’ and nearby assembly *B/F* compared to assembly *C/E* and *D*. This synaptic plasticity embeds the gradient of co-activation in the map of assemblies to outlast the effect of atomoxetine itself.

Next, we simulated the consequences of learning the ring of associations, by strengthening the excitatory (E-E) connections between neighbouring cell assemblies. Inhibitory plasticity balanced the surplus excitation by strengthening relevant inhibitory (I-E) connections between neighbouring nodes. Together, this resulted in the formation of a ring structure, with each node being connected to two other nodes via both excitatory and inhibitory connections (Figure 5B, Left). Despite strong excitatory connections between neighbouring assemblies, no significant co-activation was observed along the ring structure when externally activating one cell assembly due to the proportionally strengthened inhibitory connections.

In the computational model we then simulated the effect of learning under elevated noradrenaline (‘atomoxetine’). Unlike in the placebo model, in the atomoxetine model the associations were learned under a regime of reduced inhibitory firing. Consequently, inhibitory (I-E) connections between nodes were only minimally strengthened in response to learning. This resulted in an excitatory surplus which was not balanced by synaptic inhibition, leading to elevated excitatory plasticity. These changes manifested as a gradient of co-activation along the ring structure, in response to activation of a single cell assembly ‘*A*’ (Figure 5C). The graded co-activation occurred relative to their respective distances from assembly ‘*A*’, with greater co-activation of nearby assemblies *B/F* in contrast to *C/E* and *D*. This graded co-activation further lead to graded plasticity across cell assemblies, indicated by the changes in synaptic weights between pairs of cell assemblies which was not observed in the placebo condition (Figure 5B-C). Overall, our computational model suggests that learning during periods of globally reduced inhibition leads to synaptic modifications that embed a spread of association across memories in the underlying cognitive map.

### Elevated noradrenaline increases the spread of association in the underlying cognitive map

To empirically test the predictions of our model, we used functional Magnetic Resonance Imaging (fMRI) to measure neural representations of task stimuli and representations of the underlying memory map. Previous studies in both animals and humans demonstrate that the hippocampus represents both spatial and abstract cognitive maps of the environment (Behrens et al., 2018; O’Keefe & Nadel, 1978). Thus, in hippocampus we predicted representations that reflect distortions in the overall task structure. In addition, based on our use of an object-in-scene (bird-in-room) paradigm, we predicted neural representation of task cues in visual object and scene processing regions, namely LOC and parahippocampal cortex. Moreover, the parahippocampal cortex provides a major input-output route for information processing between sensory cortex (e.g. LOC) and the hippocampal formation (Suzuki & Amaral, 1990; Van Hoesen et al., 1979) (Figure 6A). Analyses were therefore performed across these three regions of interest: LOC, parahippocampus and hippocampus, together with the whole imaged brain volume.

**Figure 6.**
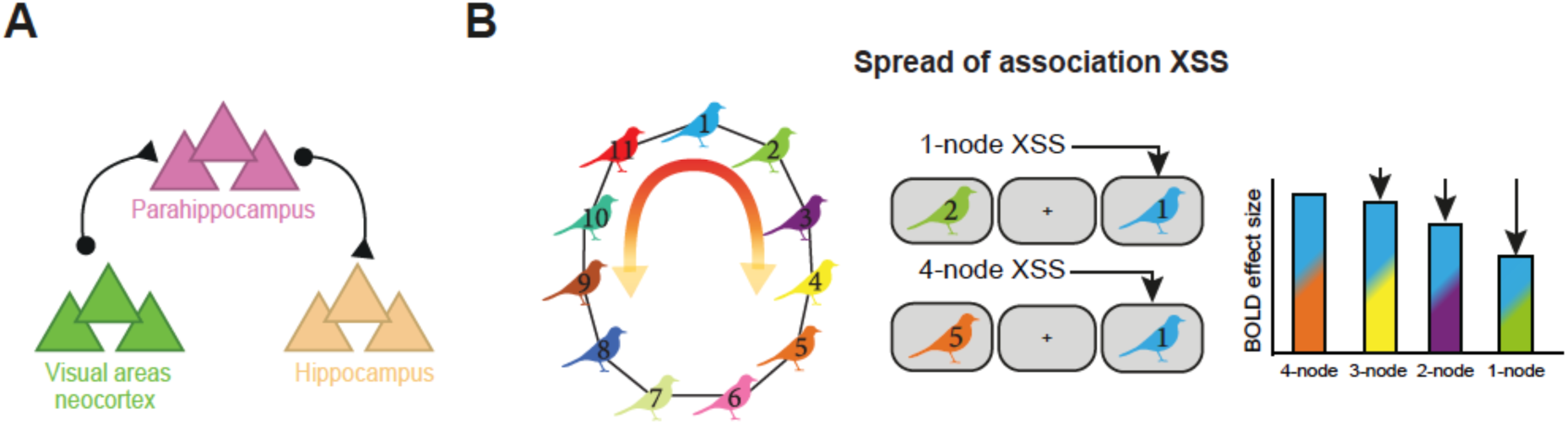
Predicted spread of association in the underlying memory map. **A** Schematic illustrating the hypothesised anatomical regions involved in representing the underlying memory map. **B** Schematic of fMRI contrast. We predicted a parametric effect in the BOLD response depending on the distance along the ring between the presented stimulus and the preceding stimulus. If a stimulus was preceded by a proximal, associated stimulus we expected to see a decrease in BOLD response (cross-stimulus suppression, XSS) compared to trials where a stimulus was preceded by a distal stimulus on the ring. This parametric regressor was used to explain variance in the BOLD signal, thus providing a measure of the spread of association in the underlying neural representation of the memory map.

To measure neural representations using fMRI we took advantage of repetition suppression, which relies on the fact that neurons show a relative suppression in their activity when presented with repeated stimuli to which they are responsive (Miller et al., 1991; Sawamura et al., 2006). While typically used with fMRI to access sub-voxel representations for single stimuli (Grill-Spector et al., 2006), “cross-stimulus” suppression can be used to index the relative co-activation or overlap between neural representations coding for two different stimuli (Barron et al., 2016). To measure cross-stimulus suppression, stimuli were presented in a pseudo-randomized order during the fMRI scan task (Figure 3A). Each stimulus was categorised post-hoc according to the node distance of the stimulus that came immediately before, as defined by the underlying ring structure (Figure 6B). For example, bird stimuli preceded by an associated bird were categorised as proximal ‘1 node’ trials (e.g. bird ‘1’ preceded by bird ‘2’), while bird stimuli preceded by birds further away on the ring structure were categorised accordingly (e.g. ‘4 nodes’: bird ‘1’ preceded by bird ‘5’) (Figure 6B). Notably, the symmetrical ring topology of the underlying memory map provided an efficient way to measure node distance from each node.

We reasoned that if noradrenaline leads to a spread of association in the underlying memory map, we should observe greater co-activity between neural representations of proximal compared to distal nodes. In the ATX group, we therefore predicted greater cross-stimulus suppression between proximal compared to distal nodes. To test this prediction, we identified brain regions where cross-stimulus suppression scales with node distance. Consistent with our prediction, in the ATX group we observed significant evidence for a spread of association across the underlying memory map in both hippocampus and parahippocampus, verified using *post-hoc* tests (Figure 7A,B, Supplementary Table 2). Importantly, no significant effect was observed in the PLC group (Supplementary Figure 8B), although no significant difference was observed between the two groups (Supplementary Figure 8C). As the response to drug was variable in the ATX group, in subsequent analyses of the underlying neural representations we took advantage of behavioural measures and our two physiological markers indicating individual differences in the response to drug.

**Figure 7.**
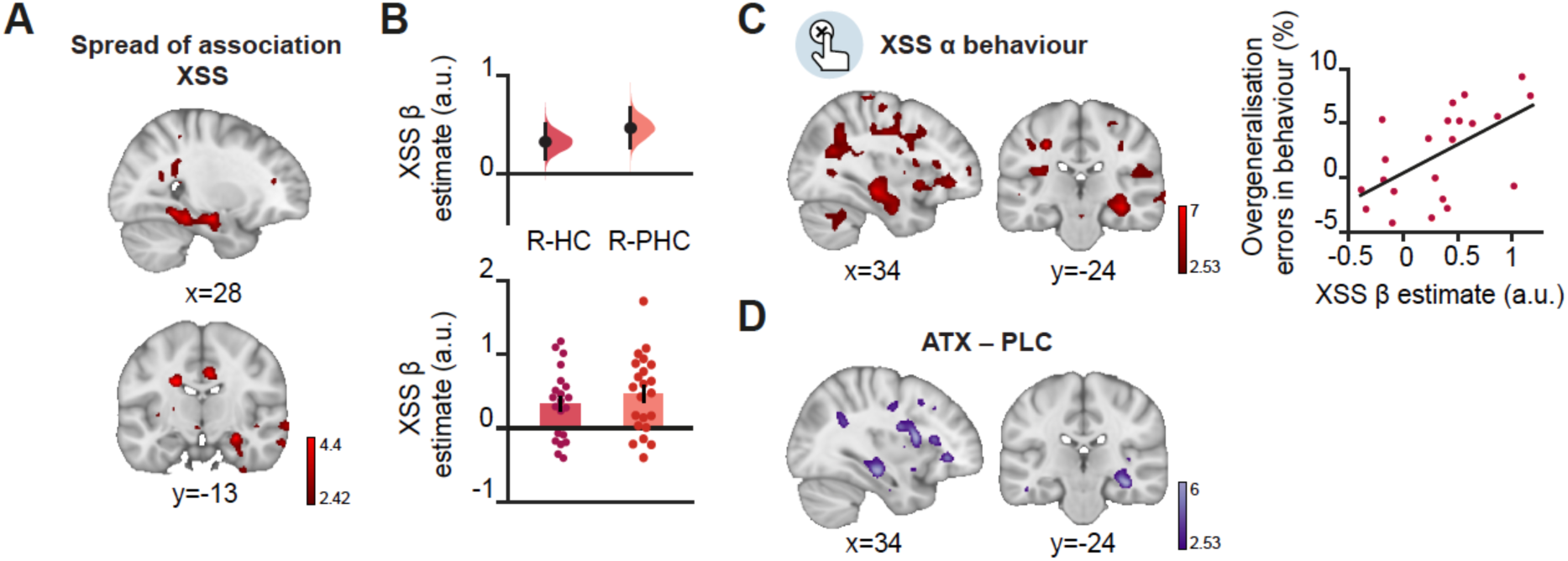
Neural spread of association in the underlying memory map predicts behavioural measure of overgeneralisation. **A** T-statistic map showing cross-stimulus suppression (XSS) for spread of association in the ATX group, with significant effect in the right hippocampus and right parahippocampal cortex (SVC with parahippocampus-hippocampus ROI: t_21_=4.56, p=0.032, MNI coordinates, Supplementary Table 2). Peaks of the effect were observed both in hippocampus and parahippocampal cortex, verified using *post hoc* tests (SVC with anatomical hippocampal ROI: t_21_=4.35, p=0.033; SVC with anatomical parahippocampal ROI: t_21_=4.56, p=0.011, MNI coordinates, Supplementary Table 2). **B** For visualisation purposes only, illustration of the effect observed in *A*, using extracted β parameter estimates from anatomical ROIs (Supplementary Figure 8A). Lower: individual parameter estimates obtained from the right hippocampus (R-HC) and right parahippocampus (R-PHC) (mean +/− SEM). Upper: Bootstrap-coupled estimation (DABEST) plots. Black dot, mean; black ticks, 95% confidence interval; filled curve, sampling error distribution. **C** Left: In ATX, the neural measure of spread of association in hippocampus and parahippocampal cortex predicted overgeneralisation errors reported in behaviour (SVC with parahippocampus-hippocampus ROI: t_20_=6.16, p=0.002, MNI coordinates, Supplementary Table 4). The peak of the effect was observed in hippocampus, verified using *post hoc* tests (SVC with hippocampal ROI: t_20_=6.16, p=0.001, Supplementary Table 4). Right: For visualisation purposes, illustrative correlation plot using extracted parameter estimates from anatomical hippocampus ROI (Supplementary Figure 8A) (Spearman correlation: r_12_ = 0.682, p=0.001). **D** There was a significant group difference (ATX vs. PLC) for the correlation between the neural spread of association (XSS) and behavioural measure of overgeneralisation errors (ATX – PLC, SVC with parahippocampus-hippocampus ROI, t_39_=4.51, p=0.011, MNI coordinates, Supplementary Table 5). The peak of the effect was observed in hippocampus, verified using *post hoc* tests (SVC with anatomical hippocampal ROI: SVC with hippocampal ROI: t_39_=4.51, p=0.007, Supplementary Table 5). T-statistic maps are thresholded at p < 0.01 uncorrected, for visualisation purposes only.

First, we asked whether the neural spread of association in the underlying memory map observed immediately after learning (Figure 7A-B) can predict overgeneralisation errors on day 5 (Figure 2F). We reasoned that participants who show stronger neural spread of association in their underlying memory map immediately after learning (Figure 6B, Figure 7A-B) should retain this knowledge and show stronger evidence for spread of association in subsequent behaviour. In the ATX group, we observed a significant positive relationship between the neural measures of spread of association in hippocampus and parahippocampal cortex and overgeneralisation errors reported in behaviour. The peak of the effect was observed in hippocampus, verified using *post hoc* tests (Supplementary Table 4). Therefore, participants who showed a greater spread of association in the underlying memory map on day 1 went on to show a greater spread of association in their behaviour on day 5 (Figure 7C). Critically, no significant effect was observed for the PLC group (Supplementary Figure 8D) and a significant difference was observed between the two groups (Figure 7D, Supplementary Table 5). Taken together, these results support the predictions from the model by suggesting that learning under elevated noradrenaline leads to a spread of association in the underlying cognitive map. The spread of association manifests as a systematic and sustained overgeneralisation in the behavioural readout of memory.

### Physiological markers of noradrenaline predict neural spread of association

These findings are observed after only a single dose of atomoxetine with variable effect across individuals. Importantly, here we measured the individual response to drug using two physiological markers of noradrenergic arousal, namely the significant increase in pupil dilation response to surprising stimuli (Figure 3B) and the significant reduction in GABAergic tone in LOC (Figure 4C). Using these two physiological markers as a measure for the response to drug, in the ATX group we asked whether individual differences in the neural spread of association in parahippocampus-hippocampus measured using fMRI can be predicted by the physiological response to drug.

Within the ATX group, we observed a significant negative relationship between GABA+ in LOC and the neural spread of association in parahippocampus-hippocampus (Figure 8A). The peak of the effect was observed in parahippocampus, verified using *post hoc* tests (Supplementary Table 6). Critically, there was no significant evidence for this relationship in the PLC group (Supplementary Figure 8E). Therefore, participants in the ATX group who learned the memory map with lower GABA+ levels in LOC showed stronger neural spread of associations in their underlying memory map. This suggests that the spread of association in the underlying memory map can be attributed to local ‘runaway excitation’, which differentially affects proximal compared to distal nodes on the ring.

**Figure 8.**
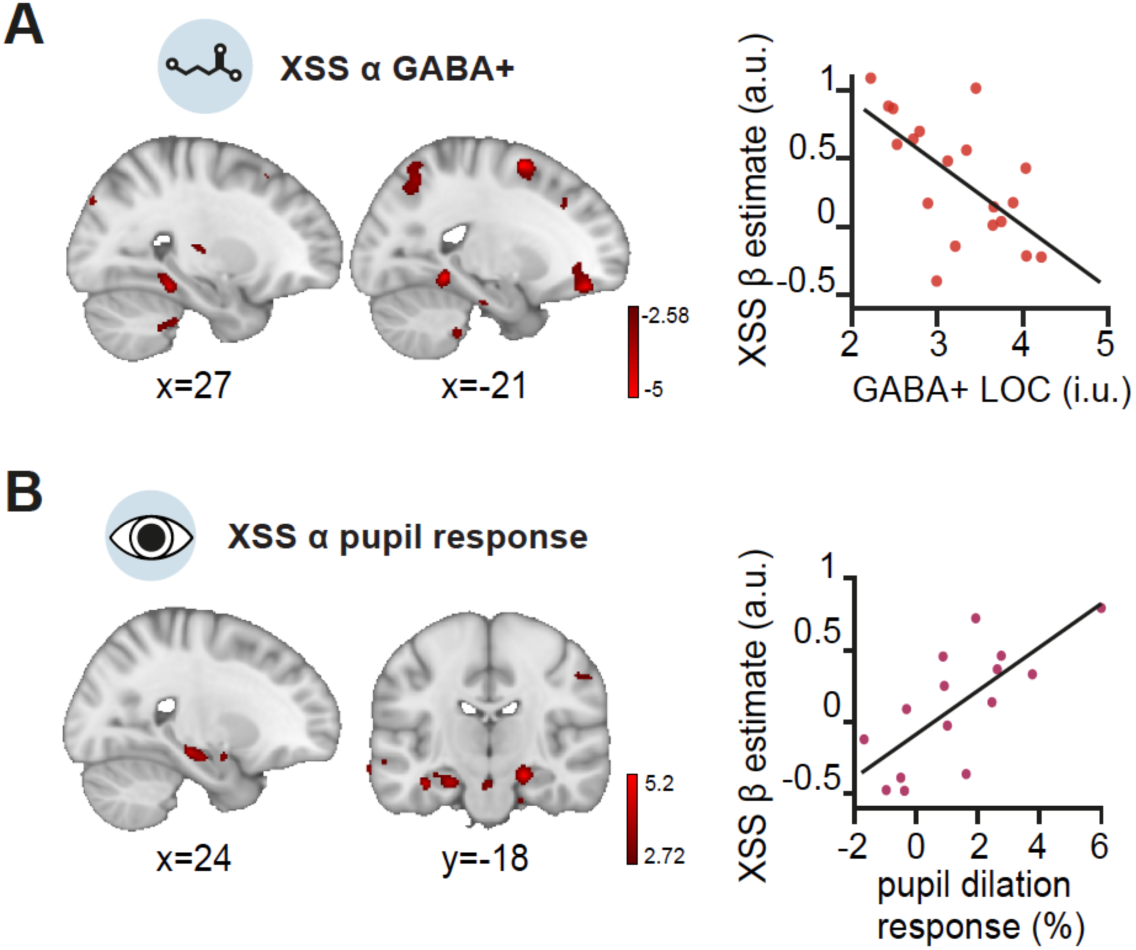
Physiological markers of reduced inhibitory tone and pupil dilation response predict neural spread of association in the parahippocampus-hippocampus. In the ATX group we used our physiological markers (GABA+ in LOC, pupil dilation response) of the response to drug to assess variance in the neural spread of association in the underlying memory map. Left: T-statistic map of the relationship. Right: illustrative correlation plot for the anatomical ROI where the effect was observed. **A** Left: The concentration of GABA+ in LOC predicted the neural measure of spread of association in hippocampus and parahippocampal cortex (SVC with parahippocampus-hippocampus ROI, t_16_=4.70, p=0.047, MNI coordinates, Supplementary Table 6). The peak of the effect was observed in parahippocampus, verified using *post hoc* tests (SVC with anatomical parahippocampal ROI, _16_=4.70, p=0.016, MNI coordinates, Supplementary Table 6). Right: For visualisation purposes, illustrative correlation plot using anatomical parahippocampal ROI (Spearman correlation: r_17_ = −0.661, p=0.003). **B** Left: We observed a positive trend between the pupil dilation response to surprising stimuli (Figure 3B) and the neural measure of spread of association in hippocampus and parahippocampal cortex (SVC with parahippocampus-hippocampus ROI, t_11_=5.15, p=0.066). A significant peak of the effect was observed in hippocampus, verified using *post hoc* tests (SVC with anatomical hippocampal ROI, t_11_=5.15, p=0.045, MNI coordinates, Supplementary Table 7). Right: For visualisation purposes, illustrative correlation plot using anatomical hippocampal ROI (Spearman correlation: r_12_ = 0.750, p=0.002). T-statistic maps are thresholded at p < 0.01 uncorrected, for visualisation purposes only.

We next asked whether our second physiological measure of response to drug, namely the pupil response to surprising stimuli, could also predict the neural spread of association in parahippocampus-hippocampus. Within the ATX group, we observed a positive trend between the pupil response and the neural spread of association in parahippocampus-hippocampus (Figure 8B). The peak of the effect was significant in hippocampus, verified using *post hoc* tests (Supplementary Table 7). Therefore, participants in the ATX group who showed a stronger pupil response also showed a stronger neural spread of association in their representation of the underlying memory map in hippocampus. No such relationship was observed in the PLC group (Supplementary Figure 8G). Taken together, these results suggest that the significant spread of association in the underlying memory map of the ATX group can be predicted by variance in the response to drug.

## Discussion

Memories are prone to distortion, but it is unclear how these distortions arise. Here, using a double-blind study design with a drug invention in humans together with computational modelling, we show that learning during a period of elevated noradrenaline alters the organisation of new memories by introducing a ‘spread of association’. This spread of association affects the underlying cognitive map by effectively increasing the size of the smoothing kernel applied to nodes within the map. The extent to which this spread of association is observed in neural readouts of the cognitive map predicts overgeneralisation errors in behaviour.

Using a multimodal approach, we combine a pharmacological manipulation with neuroimaging and behaviour to reveal a causal role for the neuromodulator noradrenaline in changing the organisation of memory. First, in the drug group who received a single dose of the noradrenergic reuptake inhibitor atomoxetine, we show that physiological markers for noradrenergic arousal are elevated. Thus, compared to placebo controls, the drug group show a larger pupil dilation response to surprising stimuli, and reduced cortical GABAergic tone. We capitalise on these physiological markers of noradrenergic arousal to assess individual differences in the response to drug. In the drug group, we observe significant spread of association across the newly learned cognitive map in the parahippocampus and hippocampus. This spread of association is predicted by physiological markers for the response to drug. In turn, we show that the spread of association in the underlying cognitive map predicts overgeneralisation errors in behaviour four days later. Taken together, these findings suggest that learning under elevated noradrenaline reduces cortical inhibition to allow for ‘runaway excitation’ across the cognitive map. This leads to a spread of association in the cognitive map that can manifest as systematic and sustained distortion in memory.

Noradrenaline has been associated with signalling salience and arousal by modulating attention and neural gain (Aston-Jones & Cohen, 2005; Bouret & Sara, 2005; Eldar et al., 2013). Noradrenaline is primarily released from locus coeruleus (LC), which projects across the entire nervous system (Berridge, 2008; Fuxe et al., 2012). Using a noradrenergic reuptake inhibitor, we demonstrate that individuals may be susceptible to memory distortions or overgeneralisation errors when learning in a state of elevated noradrenaline. In the extreme, learning in states of high noradrenergic arousal may render a memory maladaptive, as observed in Post-Traumatic Stress Disorder (PTSD). Traumatic events are typically accompanied by release of neuromodulators such as noradrenaline, and in PTSD levels of noradrenaline are higher than healthy controls and predict the severity of clinical symptoms (Kosten et al., 1987; Yehuda et al., 1998). Our findings demonstrate *how* memory distortions and systematic biases in memory can be introduced, raising an important prediction that learning under elevated noradrenaline can account for clinically significant maladaptive memories in PTSD.

While a spread of association across memories may lead to maladaptive memory distortions, our findings reveal a mechanism that carries potential for an adaptive advantage. By allowing for synthesis and integration across memories, noradrenaline appears to play two fundamental roles in the formation of a cognitive map. First, our findings show that noradrenaline provides a mechanism to bind events that occur within a given memory episode. This finding is consistent with theory and evidence to suggest that noradrenaline release demarcates event boundaries (Bouret & Sara, 2005; Clewett et al., 2020; McKenzie et al., 2024) and signals changes in the underlying cognitive map, as indexed by pupillometry data (Browning et al., 2015; Muller et al., 2019; Nassar et al., 2012). Therefore, an increase in noradrenaline at an event boundary allows for a spread of association across events that fall within a given episode. Second, noradrenaline may promote formation of a cognitive map that is more than the sum of its parts. Cognitive maps are thought to promote predictions that go beyond direct experience, to allow agents to take novel shortcuts (Gupta et al., 2010; Tolman, 1948; Wu & Foster, 2014), infer unobserved relationships (Barron et al., 2020), and generalise information across discrete experiences (Liu et al., 2019). By entering a regime of enhanced synaptic plasticity, learning under elevated noradrenaline may therefore promote representation of unobserved associations, by “joining-the-dots” between nearby nodes in the cognitive map. This mechanism may complement activity observed in the hippocampus during periods of rest and sleep (Buzsáki, 2015; Foster, 2017; Joo & Frank, 2018), where spiking sequences are generated that extend beyond direct experience (Barron et al., 2020; Dragoi & Tonegawa, 2011) to build unobserved relationships in the underlying cognitive map.

The extent to which an underlying cognitive map preserves relationships that exist in the world around us, whether observed or unobserved, may determine whether memory distortions manifest in behaviour. In humans, behavioural studies suggest that distorted memory maps provide a better explanation for behaviour than objective maps (Bellmund et al., 2020; McNamara, 1986; Warren et al., 2017). For example, measures of distance judgement, direction and map recognition in humans all suggest that cognitive maps reflect a warped version of our lived experience. Consistent with these behavioural findings, spatially tuned place cells in the hippocampus over-represent reward (Butler et al., 2019), and distortions are also reported in the spatial representations of grid cells recorded from medial entorhinal cortex (Krupic et al., 2015). Given evidence to suggest that these neuronal codes have a broader role in the organisation of memories and knowledge (Behrens et al., 2018; Constantinescu et al., 2016), these data suggest that distortions may also be observed in the mapping of more abstract stimuli and episodes. Our data provide important insight into the circumstances under which overgeneralisation errors and potential distortions can arise, demonstrating that distortions are more likely to manifest in a cognitive map when learning occurs during periods of elevated noradrenaline.

Our data provide insight into the physiological mechanisms that allow for a spread of association in the underlying cognitive map. In the drug group, we demonstrate that a single dose of atomoxetine leads to reduced cortical GABA+ in LOC. Moreover, in the drug group we observe an increase in the ratio between Glx:GABA+, suggesting that noradrenaline shifts participants’ E/I ratio along a continuum in the direction of increased cortical excitability. These findings are consistent with data showing that noradrenaline modulates the induction threshold for synaptic plasticity (Hagena et al., 2016; Lemon et al., 2009; Lisman & Grace, 2005; Tsetsenis et al., 2022) by dramatically reducing firing of inhibitory interneurons in cortex (Martins & Froemke, 2015). In cortical circuits, the effect of reducing activity in inhibitory interneurons is two-fold. First, inhibitory interneurons regulate activity of excitatory pyramidal neurons and gate the induction of excitatory synaptic plasticity (Artinian & Lacaille, 2018; Lucas & Clem, 2018). By reducing spontaneous inhibitory cell firing, noradrenaline therefore allows for a regime of enhanced excitatory synaptic plasticity (Froemke, 2015). Second, by reducing inhibitory cell firing, noradrenaline can also affect inhibitory plasticity (Hennequin et al., 2017). Inhibitory plasticity is thought to play a critical role in stabilising new learning (Barron et al., 2016; Vogels et al., 2011), by forming ‘inhibitory engrams’ (Barron et al., 2017) that protect new learning from interference (Koolschijn et al., 2019). Learning during a period of elevated noradrenaline may therefore prevent formation of inhibitory engrams, and/or render inhibitory engrams ineffective. Taken together, noradrenaline can have a profound effect on cortical plasticity, as observed when stimulating locus coeruleus, the principal source of noradrenergic modulation for the central nervous system (Martins & Froemke, 2015). Our findings further demonstrate that noradrenaline alters plasticity within a cognitive map, to change the organisation of memory with sustained behavioural consequence.

Notably, in the drug group we only observed a significant reduction in GABA+ in LOC and not in V1. The absence of an effect in V1 may be explained by significantly lower SNR in V1 compared to LOC (Supplementary Figure 7), which may reflect inhomogeneities in the static magnetic field (B0) and/or differences in physiological noise levels (Hutton et al., 2011). Alternatively, this result may be attributed to differences in the availability of noradrenaline receptors between LOC and V1, given known variation in noradrenergic receptor and fibre density across sensory cortex and across different cortical sub-regions (Agster et al., 2013; Chandler et al., 2014; Chandler & Waterhouse, 2012; Kebschull et al., 2016; Lewis & Morrison, 1989). In addition, depending on the cortical region, atomoxetine may differentially affect other neuromodulators. For example, in the rat prefrontal cortex, atomoxetine increases extracellular levels of noradrenaline as well as dopamine (Bymaster et al., 2002; Swanson et al., 2006), where dopamine reuptake is regulated by noradrenaline transporters (Bymaster et al., 2002). Atomoxetine has also been shown to increase extracellular levels of acetylcholine in the rodent cortex and hippocampus, and these procholinergic effects are hypothesised to be related to (working) memory improvements associated with atomoxetine (Callahan et al., 2019; Tzavara et al., 2006). Critically, in our data the significant reduction in GABA+ in LOC, together with changes in pupil response, provide a measure of the response to drug which significantly predicts the extent to which the spread of association is observed in the underlying cognitive map. Thus, by quantifying the response to drug using known physiological markers of noradrenergic arousal, we can account for variance in the response of any given individual or brain region to atomoxetine.

In summary, here we show that an increase of the neuromodulator noradrenaline during learning induces a ‘spread of association’ in the underlying hippocampal cognitive map. The neural spread of association in hippocampus can be explained by physiological markers of noradrenergic arousal, namely a reduction in GABAergic tone and an increased pupil dilation response. The neural spread of association effect further predicts systemic and sustained overgeneralisation errors in the behavioural read-out of memory. These findings provide mechanistic insight into how the neuromodulator noradrenaline shapes the organisation of information within a cognitive map, by increasing the width of the ‘smoothing kernel’ for plasticity across the cognitive map.

## Methods

### Participants and double blinding

44 healthy volunteers participated in the study. We implemented a double-blind, randomized, placebo-controlled study design. Participants were randomly assigned to one of two groups which were matched by sex: one group who received a single dose of 10mg atomoxetine (ATX group, n=22, mean age: 23.9 +/− 5.23 yrs, 11 women) and one group who received placebo (PLC group, n=22, mean age: 25.2 +/− 4.80 yrs, 11 women). All experiments were approved by the University of Oxford ethics committee (reference number R60579/RE004). All participants gave informed written consent.

Randomisation was performed by a researcher not involved in recruitment and data collection (LC). The drugs were administered by an experimenter (RK), who was blind to the drug condition. At the end of the final testing day, participants and experimenter indicated whether they thought the participant had received atomoxetine or placebo.

The effectiveness of the double blinding procedure was assessed using the Bang’s Blinding Index (BI). A score of 1 indicates that all responses were correct and complete unblinding is inferred; a score of −1 indicates that all responses were incorrect and complete blinding is inferred or unblinding in the opposite direction (e.g. opposite guessing); a score of 0 indicates that half the guesses were correct and half of the guesses were incorrect, inferring successful blinding with random guessing. Scores between −0.2 and 0.2 indicate that blinding was successful.

### Behavioural training

On the first day of the experiment, all participants (n=44) were given an over-encapsulated capsule, which contained either a single dose of 10mg atomoxetine (ATX group) or an empty capsule (PLC group). Participants were then required to wait for 90 minutes (Figure 1G) to allow time for the drug to take effect (Sauer et al., 2005).

Following the waiting period, participants learned 11 pairwise associations between bird stimuli. Each pair of birds was presented in a particular context, where the context constituted a living room interior (Figure 1B,C). The bird stimuli were always presented in the same location within each particular context (Supplementary Figure 2). Together, the context dependent pairwise associations formed a symmetrical ring structure (Figure 1A). This ring structure provided an efficient way to later measure representational overlap between proximal and distal associative links as each bird could be used as a control for an association on the other side of the ring (see ***fMRI scan task***). Participants were not made aware of the underlying ring structure.

To learn the bird-bird associations, participants performed a 3-alternative forced choice task with feedback (‘learning task’, Figure 1D). Every trial, participants were shown a background stimulus with one of the bird stimuli in place. Choice stimuli could be displayed upon pressing the buttons ‘b’,’n’, or ‘m’ on the keyboard and participants confirmed their choice by pressing one of these three buttons together with the spacebar. There was no response deadline. Feedback was subsequently presented for 1.5 s, after which the background with the correct stimulus pairing was shown until the participant again pressed spacebar to proceed to the next trial. Participants were required to score >90% correct on the 3-alternative forced choice learning task for two consecutive blocks. Upon reaching this criterion, participants completed an inference test (Supplementary Figure 3A) which involved participants indicating which bird stimulus shared an association with the probe bird (e.g. probe bird: bird ‘1’; inferred choice: bird ‘3’) in the absence of contextual cues or feedback, again using a 3-alternative forced choice test. Importantly, throughout these training tasks the bird-bird associations were made explicit while the bird-sofa relationships remained implicit.

Four days later, on Day 5, participants completed a battery of tests to assess memory for explicitly and implicitly learned associations. The battery of tests included the following: (1) ‘Explicit Memory Test’, where a 3-alternative forced choice task was used to test memory for bird-bird associations in the absence of contextual cues or feedback (Figure 2A); (2) a repeat of the ‘Inference Test’, as encountered on day 1 and described above (Supplementary Figure 3A); (3) ‘Implicit Memory Test’, where a 6-alternative forced choice task was used to test memory for bird-context associations in the absence of feedback (Figure 2B). In the Implicit Memory Test images of sofas taken from the background interiors were used as contextual cues, rather than the full image of the living room interior. Importantly, throughout the task the bird stimuli were always presented in the same location within their contextual interior but never on or proximal to the sofas.

For each memory test, average accuracy was estimated for each participant. On the Implicit Memory Test, as overall memory accuracy was close to chance, we analysed the error trials to estimate a behavioural ‘spread of association’ between stimuli, i.e. the tendency of participants to overgeneralise by associating a bird stimulus with an incorrect contextual cue that was proximal rather than distal along the ring. For example, when prompted with bird 1, an overgeneralisation error involved selecting, for example, sofa B rather than sofa A, where sofa B was presented with bird 2 and bird 3 making it an incorrect but *proximal* choice. As such, overgeneralisation errors were defined as the percentage of trials where participants erroneously linked a bird stimulus to a proximal (neighbouring) contextual cue in the underlying ring structure (Figure 2E, e.g. selecting sofa B when prompted with bird 1), minus the percentage of trials where any other error was made (e.g. selecting sofa E when prompted with bird 1).

### MRS data collection and analysis

MRS spectra were acquired using a MEscher-GArwood Point RESolved Spectroscopy (MEGA-PRESS) sequence in two 2 x 2 x 2 cm3 VOIs, with TE = 68 ms, TR = 1.5 s, flip angle 90 deg, ON editing pulse 1.9 ppm, OFF editing pulse 7.5 ppm. Unsuppressed water signal was collected for internal water referencing. The first volume of interest (VOI) was centered bilaterally on the calcarine sulcus in visual cortex (V1). The second VOI was positioned in right LOC. For each measurement 320 spectra were acquired (160 edit-ON and 160 edit-OFF), resulting in an acquisition time of about 8 min for each VOI. B0 shimming was performed using a GRE-SHIM (FOV =216 mm2; TR= 482 ms; TE1/2 = 4.92/7.38 ms; slice thickness 2 mm; flip angle 46 deg; slices 49; scan time 105 s).

Glx and GABA+ concentrations were quantified using Gannet version 3.1.4 (Edden et al., 2014), with water used as a reference. Gannet’s standard preprocessing pipeline was used, which includes frequency and phase correction by spectral registration and line broadening. Individual ON and OFF subspectra were then averaged and edited spectra were generated by subtracting the averaged edit-OFF spectra from edit-ON spectra. Notably, the editing approach not only targets GABA but also other macromolecules at 3ppm, therefore the concentrations of GABA+ (GABA and macromolecules) are reported. Furthermore, the editing approach also targets nearby glutamate/glutamine resonances, providing an estimate of Glx concentration. However, it should be noted that the MEGA-PRESS uses parameters optimised for editing GABA and not glutamate. Therefore, compared to quantifying Glx using a PRESS sequence, test-retest reliability of the Glx is likely compromised when estimated with MEGA-PRESS (Van Veenendaal et al., 2018; Maddock et al., 2018). Metabolite concentrations were quantified in pseudo-absolute molality units (approximating moles of GABA per kg of solute water) and relaxation and tissue-corrected (Gasparovic et al. 2006). Grey matter, white matter and CSF tissue fractions for determining tissue-corrected concentrations were obtained for both using SPM12.

Reliable model fits were achieved for 38 out of 41 V1 acquisitions (ATX: 20, PLC: 18) and for 37 out of 43 LOC acquisitions (ATX: 19, PLC: 18). In the remaining acquisitions GABA+ was unreliable or inestimable due to lipid contamination or low SNR. Data quality was quantified using the signal-to-noise ratio (SNR) and full-width-at-half-max (FWHM) of N-acetylaspartate (NAA) and fit error of the GABA+ peak provided by Gannet (Supplementary Figure 6). The average GABA+ and Glx concentrations for the ATX and PLC groups were compared using two-tailed permutation tests. A null distribution of the expected difference was generated by permuting group labels (ATX, PLC) 10,000 times using MATLAB’s random number generator.

### fMRI scan task

During the fMRI scan, participant viewed the 11 bird stimuli encountered during learning, across 4 scan blocks. During 2 of these scan blocks, bird stimuli were interleaved with the 11 sofa contextual stimuli. Stimuli were presented via a computer monitor inside the scanner bore. On each trial, stimuli were presented for 1000 ms (Figure 3A). The inter-trial interval was selected from a truncated gamma distribution with a mean of 2.3 s, minimum of 1.3 s and maximum of 14 s. To control for potential confounding effects of expectation suppression (Summerfield et al., 2008), all possible transitions between non-repeating stimuli were presented once in each block, in a fully randomized order. Participants performed a task incidental to the contrast of interest which involved identifying whether the presented stimuli were familiar or ‘‘oddball”. Oddball stimuli were distorted or differently coloured versions of the stimuli encountered during training, and were randomly inserted into 9% of trials. Participants were instructed to press a button on an MR compatible button box using their right index finger when they identified “oddball” stimuli, but not if the stimulus was familiar. Participants received a monetary reward for each correct oddball identification. No feedback was given other than the total monetary reward after each block.

### Pupillometry acquisition, preprocessing, and analysis

During the fMRI scan task (see ***fMRI scan task***) pupillometry data was collected. We measured pupil size using an EyeLink eyetracker (SR Research) sampled at 500 Hz. Pupillometry data was preprocessed using a standardised processing pipeline. Briefly, data was smoothed using a 120ms Gaussian kernel, blinks were identified and removed by interpolating the pupillometry data across the blink period. Data were epoched around presentation of the visual stimuli. Trials on which the pupil measurement was lost for a period of >1000ms were removed from the analysis. Participants were excluded from subsequent analysis if more than 50% of trials had been removed (n=10 participants). Overall, pupillometry data from 34 out of 44 participants was included (ATX: 15, PLC: 19), with an average of 672.50 ± 17.69 ‘regular’ trials and 58.82 ± 1.36 ‘oddball’ trials (mean ± SEM).

To normalise pupil size for each scan block we expressed pupil size in terms of the percentage difference from the average pupil size across the block. To account for changes in pupil size over the course of the fMRI scan task, each epoched pupil response was baseline corrected. The baseline was defined as the mean of the normalised pupil size in the 250 ms window prior to stimulus presentation. To assess the effect of surprise on the pupil response, for each participant we computed the difference between the epoched pupil response for ‘oddball’ vs ‘regular’ stimuli. To assess the difference in pupil response between the ATX and PLC groups, we used a cluster-based significance approach where we permuted subject labels x10,000 for each timepoint, to define the size threshold for a statistically significant cluster (p<0.05) (Hunt et al., 2013; Hauser et al., 2015).

### fMRI imaging protocol

The fMRI scan task was performed inside a 3 T Magnetom Prisma (Siemens) using a 32-channel head and neck coil (Nova Medical Inc, Wilmington, MA) at the Wellcome Centre for Integrative Neuroimaging (University of Oxford). fMRI data was acquired using a multiband echo-planar imaging (EPI) sequence, with multiband factor 3, TR = 1.235 s, TE = 20 ms, flip angle = 65 degrees, field of view = 216 mm, and a voxel resolution of 2×2×2 mm^3^. All scans were of axial orientation angled to the long-axis of the hippocampus, covering the whole brain. For each participant, a T1-weighted structural image was acquired to correct for geometric distortion and perform co-registration between EPIs, consisting of 192 1 mm axial slices, in-plane resolution of 1×1 mm^2^, TR= 1.9 s, TE = 3.97 ms, and field of view = 192 mm. A field map with dual echo-time images was also acquired (TE1 = 4.92 ms, TE2 = 7.38 ms, whole-brain coverage, voxel size 2×2×2 mm^3^). Physiological measures (cardiac pulse and thoracic movement) were collected during the scan but not used in analysis. Cardiac pulse was recorded using an MRI-compatible pulse oximeter. Thoracic movement was monitored using a custom-made pneumatic belt positioned around the abdomen.

### fMRI preprocessing and GLMs

Preprocessing of fMRI data was carried out in SPM12 (https://www.fil.ion.ucl.ac.uk/spm/). The dataset of one participant was excluded due to excessive motion. For the 43 participants included in the fMRI analysis, images were corrected for signal bias, realigned to the first volume of the block, corrected for distortion using fieldmaps, normalised to a standard EPI template and smoothed using a 5 mm full-width at half maximum Gaussian kernel. To remove low frequency noise from the preprocessed data, a high-pass filter was applied to the data using SPM12’s default settings. For each participant and for each scan block presenting bird stimuli, the resulting fMRI data was analysed in an event-related manner using a General Linear Model (GLM). Explanatory variables used a delta function to indicate the onset of a stimulus and were then convolved with the hemodynamic response function. In addition to the explanatory variables (EVs) of interest (described below), six additional scan-to-scan motion parameters produced during realignment were included in the GLM as nuisance regressors to account for motion-related artefacts in each task block.

In the GLM, 14 EVs were included per scan block. The first EV was the parametric regressor corresponding to the link distance (nodes) between the presented stimulus and the preceding stimulus measured along the ring structure. The parametric regressor was according to an exponential function (1–2^[link distance]) to account for non-linearity in link distance encoding, and was subsequently z-scored. The next 11 EVs accounted for each of the different bird stimuli (bird 1-11). The remaining 2 EVs accounted for detected and undetected ‘oddball’ trials, respectively. All EVs were then convolved with the hemodynamic response function. The contrast of interest was the parametric regressor of link distance (Figure 6B), contrasted against baseline.

### fMRI analysis and statistics

The contrast of interest was entered into second-level random effects ‘group’ analyses to assess the main effect of link distance, and to assess the effect of 3 covariates: GABA+ in LOC, pupil response, and behaviour. Second-level analyses were performed for each group separately (ATX, PLC), and for all participants combined. We set the cluster-defining threshold to p < 0.01 uncorrected before using whole-brain family wise error (FWE) to correct for multiple comparisons, with the significance level defined as p < 0.05. To assess changes in BOLD in regions of interest where we predicted representation of task stimuli and a ‘spread of association’ across memory (Figure 6B), we performed small volume correction for multiple comparisons, with peak-level FWE correction at p<0.05. For the small volume correction in hippocampus and parahippocampus we used bilateral anatomical masks segmented from several representative T1 images normalised to the MNI152 template (Supplementary Figure 8A,B). For small volume correction in LOC we used a 10mm radius sphere centered on the group-average MRS voxel (Supplementary Figure 7C).

To assess the relationship between fMRI cross-stimulus suppression and each of the 3 covariates of interest (GABA+ in LOC, pupil response, and behaviour), we ran the second-level random effects model with the inclusion of each covariate at the group level. For the GLM with GABA+, an additional covariate was included to account for differences in the concentration of Glx in LOC. To assess the relationship between pupil response and fMRI, a pupil response covariate was defined as the average difference in pupil response to ‘oddball’ stimuli versus regular stimuli, across the time period where we observed a group difference between ATX and PLC (Figure 3B). Covariates for the concentration of GABA+ and Glx in LOC were included to assess variance predicted by the pupil response, over and above variance explained by changes in GABA+ and Glx. Finally, to assess the relationship between fMRI cross-stimulus and the behaviour, a ‘spread of association’ effect (Figure 2F), we included a covariate for the behavioural measure of overgeneralisation errors from the implicit memory test (Figure 2F).

For visualisation purposes only, we used a region of interest (ROI) approach to visualise the relationships between fMRI cross-stimulus suppression and GABA+ in LOC, pupil response, and behaviour. To this end, we used anatomical ROIs for right parahippocampus and right hippocampus to extract beta parameter estimates for the parametric link distance regressor, and computed the Spearman rank correlation for each relationship (Figure 7A, Figure 8A–B).

### Recurrent spiking neural network model

#### Neuron Model

The postsynaptic neurons were described by point leaky integrate-and-fire neurons with after-hyperpolarisation (AHP) current, whose membrane potential *V*(*t*) evolved according to:

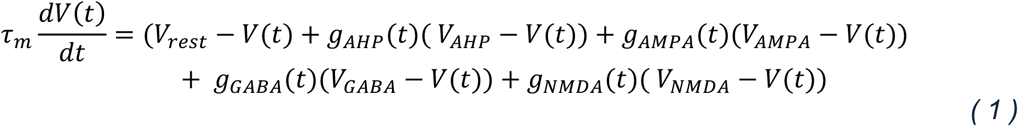

where *τ*_*m*_ = 30 *ms* is the membrane time constant and *V*_*rest*_ = −65 *mV* is the resting membrane potential; *V*_*AMPA*_ = *V*_*NMDA*_ = 0 *mV* and *V*_*GABA*_ = *V*_*AHP*_ = −80 *mV* are the reversal potentials; *g*_*AMPA*_, *g*_*NMDA*_, and *g*_*GABA*_ are the synaptic conductances for their respective channels, and *g*_*AHP*_ is the conductance of the AHP channel. The neuron fires when its membrane potential crosses a fixed threshold of *V*_*th*_ = −50 *mV* below, after which it returns to its resting potential where it remains clamped for 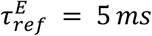 and 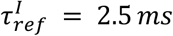 of refractory period in excitatory and inhibitory neurons, respectively.

#### Synapse Model

The synapses were conductance-based. The AHP conductance follows the equation:

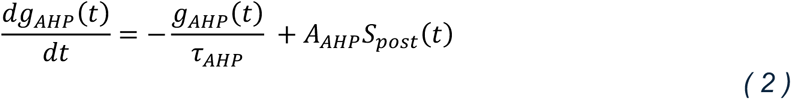

Where *τ*_*AHP*_ = 100 *ms* is the characteristic time of the AHP channel, and *A*_*AHP*_ = 5*nS* is the amplitude of increase in conductance due to a single postsynaptic spike. 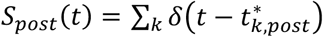 is the postsynaptic spike train, where 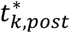 is the time of the *k*^*th*^ spike of the postsynaptic neuron, and *δ*(.) is the Dirac’s delta.

The synaptic conductance for *X* (where *X* can be *AMPA*, *GABA*, or *NMDA*) obeys the following equation:

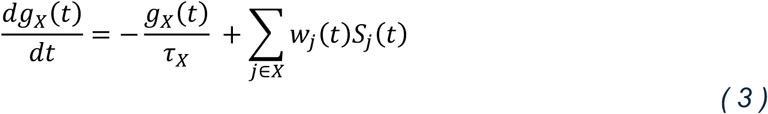

Here, the characteristic times are *τ*_*AMPA*_ = 5 *ms*, *τ*_*GABA*_ = 10 *ms*, *τ*_*NMDA*_ = 150 *ms*. 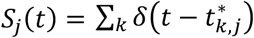 represents the presynaptic spike train, where 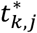 is the time of the *k*^*th*^ spike of neuron *j*.

#### Plasticity Model

The plasticity model employed in these simulations was the so-called “codependent” plasticity model (Agnes & Vogels, 2024). Codependent synaptic plasticity rules depend on both spike times and synaptic currents. The codependent plasticity function *ϕ* of a synapse is represented as:

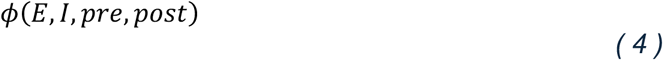

where *E* and *I* are the excitatory and inhibitory currents due to neighbourhood synaptic activity (including the synapse’s own activity), and *pre*, *post* are the spike timings of the pre- and postsynaptic neurons, respectively. *E* represents the calcium influx through N-methyl D-aspartate (NMDA) channels to model excitatory currents and *I* represents the chloride influx through gamma-aminobutyric acid (GABA) channels to model inhibitory currents. These currents are defined as follows:

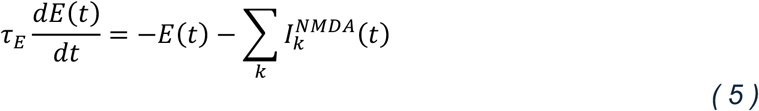

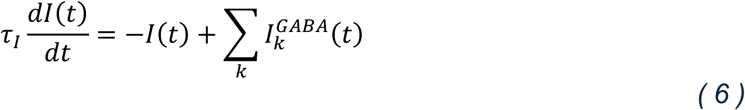

where *τ*_*E*_ = 10 *ms* and *τ*_*I*_ = 100 *ms* are the characteristic time constants for *E* and *I*, respectively. 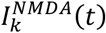 and 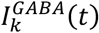 are the currents flowing through the NMDA and GABA channels for the *k*^*th*^ excitatory and inhibitory synapses, respectively.

The learning rule of the *j*^*th*^ excitatory synapse onto the postsynaptic neuron is defined as:

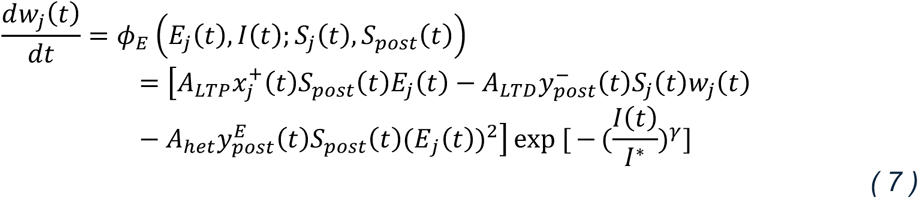

where *A*_*LTP*_ = 3 × 10^-4^ and *A*_*LTD*_ = 3 × 10^-5^ are the learning rates of Hebbian long-term potentiation and long-term depression, respectively, and *A*_*het*_ = 1.5 × 10^-8^ is the learning rate of the heterosynaptic plasticity, which depends quadratically on the neighbouring excitatory currents. *I*^∗^ = 200 and *γ* = 3 represent the level and shape of the control of inhibitory activity over changes in excitatory weights. *S_j_*(*t*) and *S*_*post*_(*t*) are the pre- and postsynaptic spike trains, as described earlier. 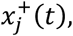 representing the trace of the presynaptic spike train, follows:

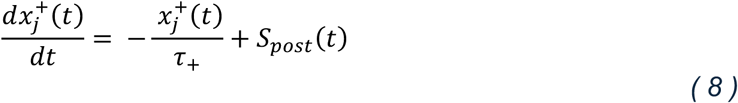

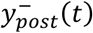 and 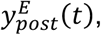 representing the traces of the postsynaptic spike trains with different time scales, follow:

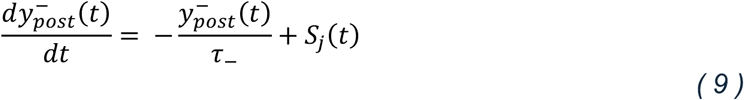

and

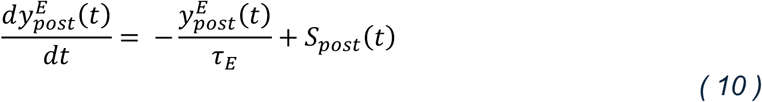

where *τ*_+_ = 16.8*ms*, *τ*_−_ = 33.7*ms*, and *τ*_*E*_ = 100*ms*.

The inhibitory learning rule of the *j*^*th*^ inhibitory synapse onto the postsynaptic neuron is defined as:

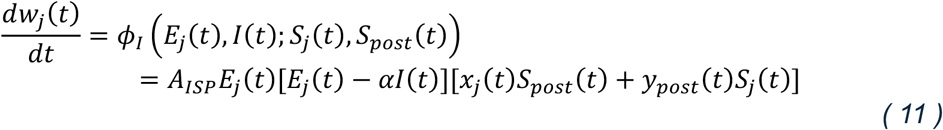

where *A*_*ISP*_ = 1 × 10^-3^ is the codependent inhibitory plasticity learning rate, *α* is the excitatory-inhibitory balance point, such that 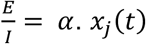 and *y*_*post*_(*t*) are the synaptic traces of the pre- and postsynaptic spike trains, respectively. They follow:

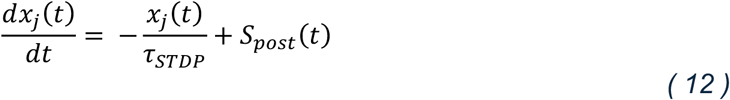

and

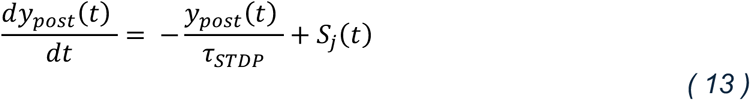

where *τ*_*STDP*_ = 20*ms*.

#### Network and Simulation

In the neural network model, we simulated a recurrent neural network of 1000 excitatory and 250 inhibitory neurons. E-E and I-E connections were plastic, while E-I and I-I connections were static. Connectivity was 5% between and across all neuron groups. Initial weights for the different synapse types were *w*_*EE*_ = 0.25 *nS*, *w*_*EI*_ = 0.35 *nS*, *w*_*IE*_ = 0.31 *nS*, *w*_*II*_ = 0.31 *nS*. Background network activity was maintained by connecting a group of 100 Poisson neurons firing at 1Hz to the excitatory neuron group with a weight of *w*_*input*_ = 0.1 *nS* and connection probability of 1.

#### Cell Assembly Formation

The cell assemblies A-F were formed by connecting six independent Poisson groups (representing external sensory stimuli) of 100 neurons each, firing at 2.5 Hz, to six groups of 100 excitatory neurons each. The synapses between the Poisson neurons and the excitatory neurons had a weight of 0.1 ± 0.01 *nS*. This resulted in an increase in network excitation, which was subsequently balanced by strengthening *w*_*IE*_I-E connections between groups of 25 inhibitory neurons to each of the formed excitatory cell assemblies to *w*_*IE*_ = 0.9 ± 0.05 *nS*. Hence, each assembly A-F was composed of 100 excitatory neurons and 25 inhibitory neurons.

#### Inter-nodal Associations

In both the placebo and atomoxetine neural network simulations, excitatory connections between formed nodes were strengthened by setting E-E connections between them to *w*_*EE*_ = 0.43 ± 0.02 *nS*. However, in the atomoxetine simulations, these E-E connections were strengthened under decreased inhibitory firing, which was achieved by setting *w*_*EI*_to only 97% of its value, or *w*_*EI*_ = 0.3395 *nS*.

In the placebo simulations, this E-E strengthening was followed by setting inter-nodal I-E connections to *w*_*IE*_ = 0.9 ± 0.05 *nS*. In the drug case, these connections were only strengthened to *w*_*IE*_ = 0.7 ± 0.05 *nS*, which was the minimum I-E strengthening required to prevent runaway excitation due to E-E potentiation triggered by the inter-nodal excitatory associations.

#### Driving Activity of Assembly ‘A’

To test the degree of coactivation, the activity in assembly ‘*A*’ was driven externally for 3.3s during the period marked “stimulus” (Figure 5, centre plots) by increasing the weight of the relevant Poisson group representing the sensory input to 0.5 *nS*.

## Data and Software availability

All data generated and analysed during this study are included in the manuscript and supporting files. Upon publication the group-level data and code used in this study are available via the MRC BNDU Data Sharing Platform. The data will be made available via https://data.mrc.ox.ac.uk/data-set/fmri-noradrenaline-memory. The code will be made available via https://data.mrc.ox.ac.uk/data-set/fmri-noradrenaline-memory-code.

The following dataset was generated:

- fMRI data
- MRS data
- Pupillometry data
- Behavioural data
- code

### Acknowledgements

We would like to thank Liliana Capitão for her advice and support with setting up the double-blinding procedure, and Chamith Halahakoon, Phil Cowen, Angharad De Cates, Beata Godlewska, Riccardo De Giorgi, Katherine Smith and Edoardo Ostinelli for enabling this study by providing medical cover. We would like to thank Douglas F. Tomé and Everton J. Agnes for their guidance and advice with earlier versions of the neural network model. We thank Leonie Glitz and Valentina Mancini for comments on an earlier version of the manuscript. R.S.K. was supported by an EPSRC/MRC-funded studentship (EP/L016052/1). P.P. was supported by the Cambridge Trust, Trinity Henry Barlow Scholarship and Trinity Hall Brockhouse Scholarship. H.C.B. is supported by a UKRI Future Leaders Fellowship (MR/W008939/1) and the Wellcome Institutional Strategic Support Fund. H.C.B. and J.X.O. are supported by the Medical Research Council (MR/W01971X/1). The study was supported by the NIHR Oxford Health Biomedical Research Centre. The views expressed are those of the authors and not necessarily those of the NHS, the NIHR or the Department of Health. The Wellcome Centre for Integrative Neuroimaging is supported by core funding from the Wellcome Trust (203139/Z/16/Z).

## Author contributions

All authors contributed to the preparation of the manuscript. R.S.K., J.X.O. and H.C.B designed the study. L.C. and R.S.K prepared and administered the double-blind procedure. M.B. provided clinical support for the study. R.S.K. acquired the data. W.C. and R.S.K. developed the MRS sequence and analysed the MRS data. R.S.K. analysed the behavioural data, pupillometry data and fMRI data with supervision from J.X.O. and H.C.B. X.P. assisted with the fMRI analyses. P.P. generated all neural network simulations with supervision from T.P.V. R.S.K., P.P. and H.C.B. prepared the figures.

## Supplementary Materials

## Supplementary Figures

**Supplementary Figure 1.**
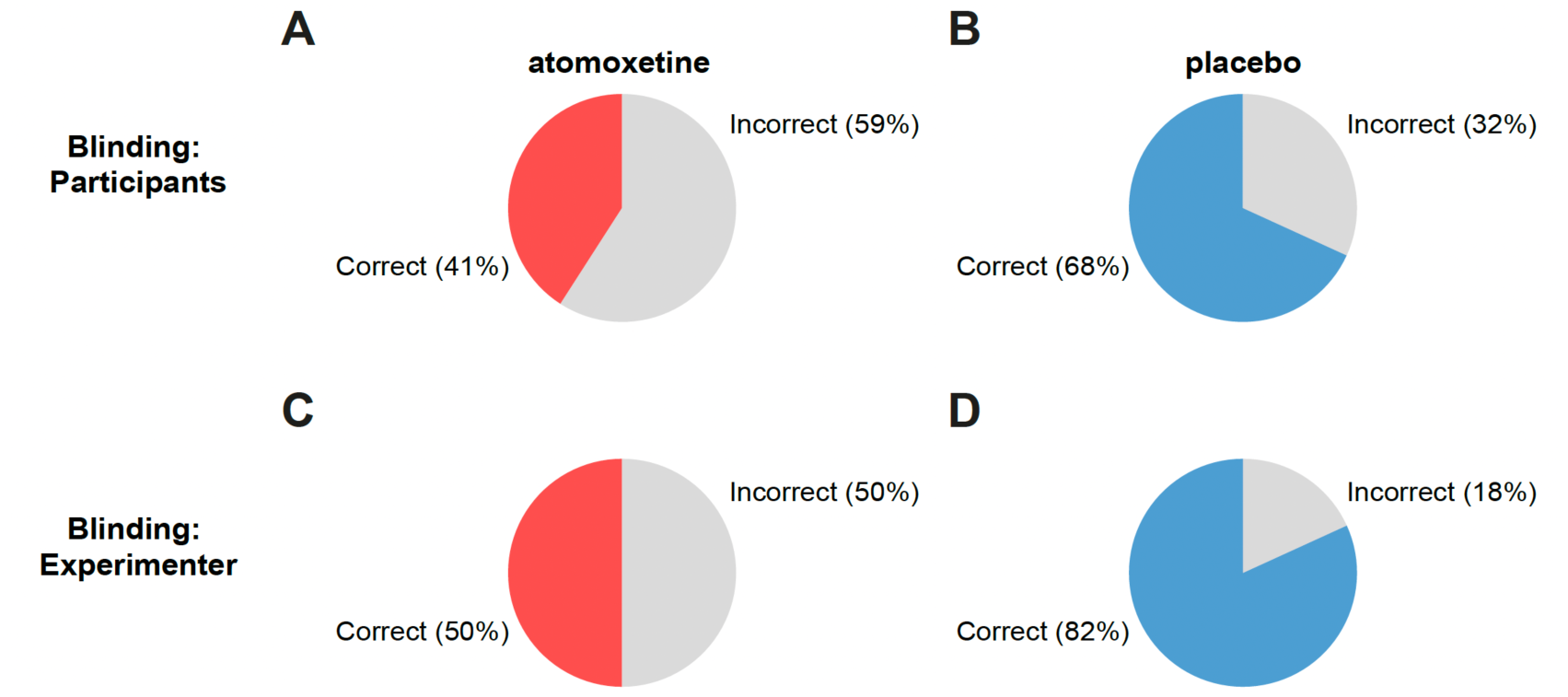
Blinding data for participants and experimenters. To assess the effectiveness of the double-blinding procedure (see also Supplementary Table 1), after completion of the first study day the participants and the experimenter were asked whether they thought they/the participant received drug or placebo. **A** In the atomoxetine group, 41% of participants correctly guessed that they had received atomoxetine. **B** In the placebo group, 68% of participants correctly guessed that they had received placebo. **C** The experimenter correctly guessed for 50% of participants that they had received atomoxetine. **D** The experimenter correctly guessed for 82% of participants that they had received placebo.

**Supplementary Figure 2.**
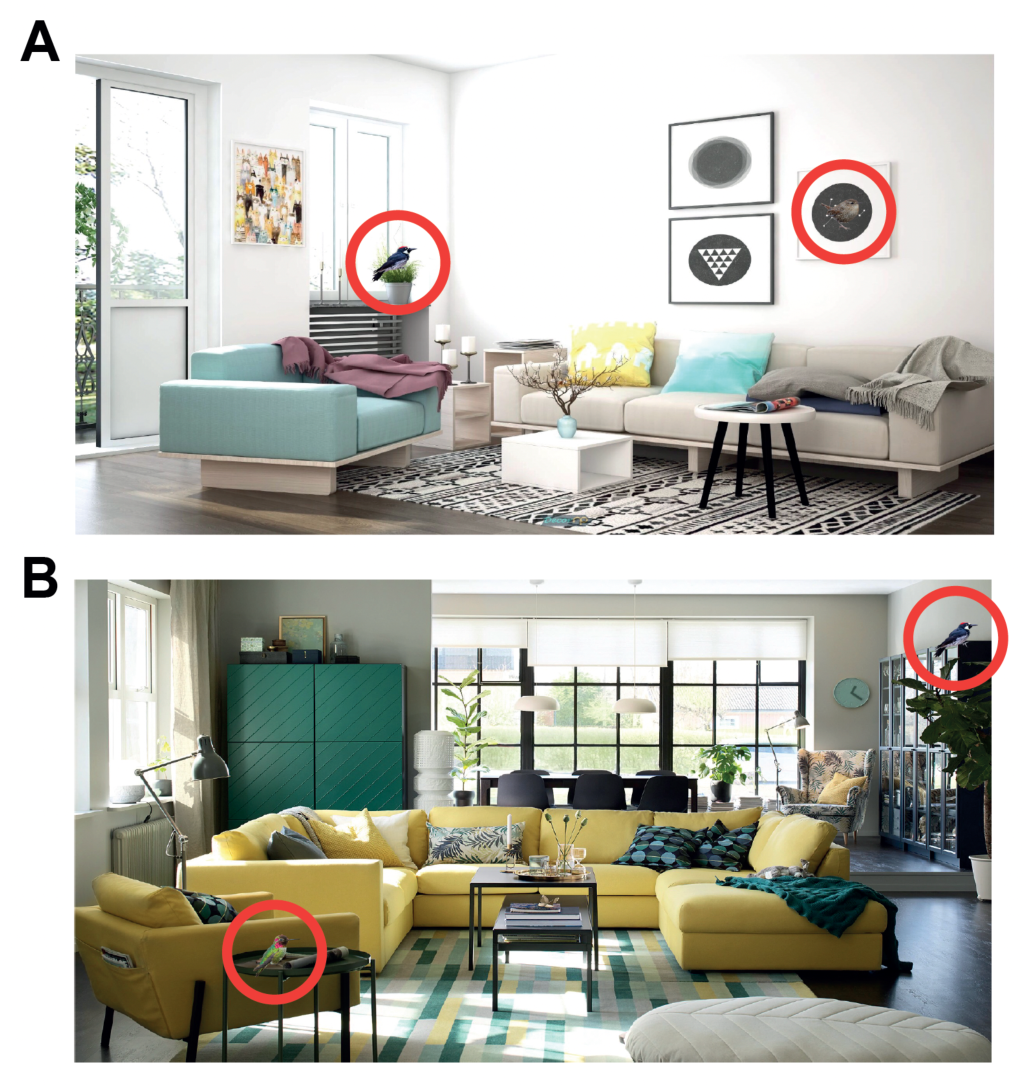
Example trials from learning task. **A-B** Two example bird pairs situated within their contextual cues. Red circles indicate the location of the bird stimuli.

**Supplementary Figure 3.**
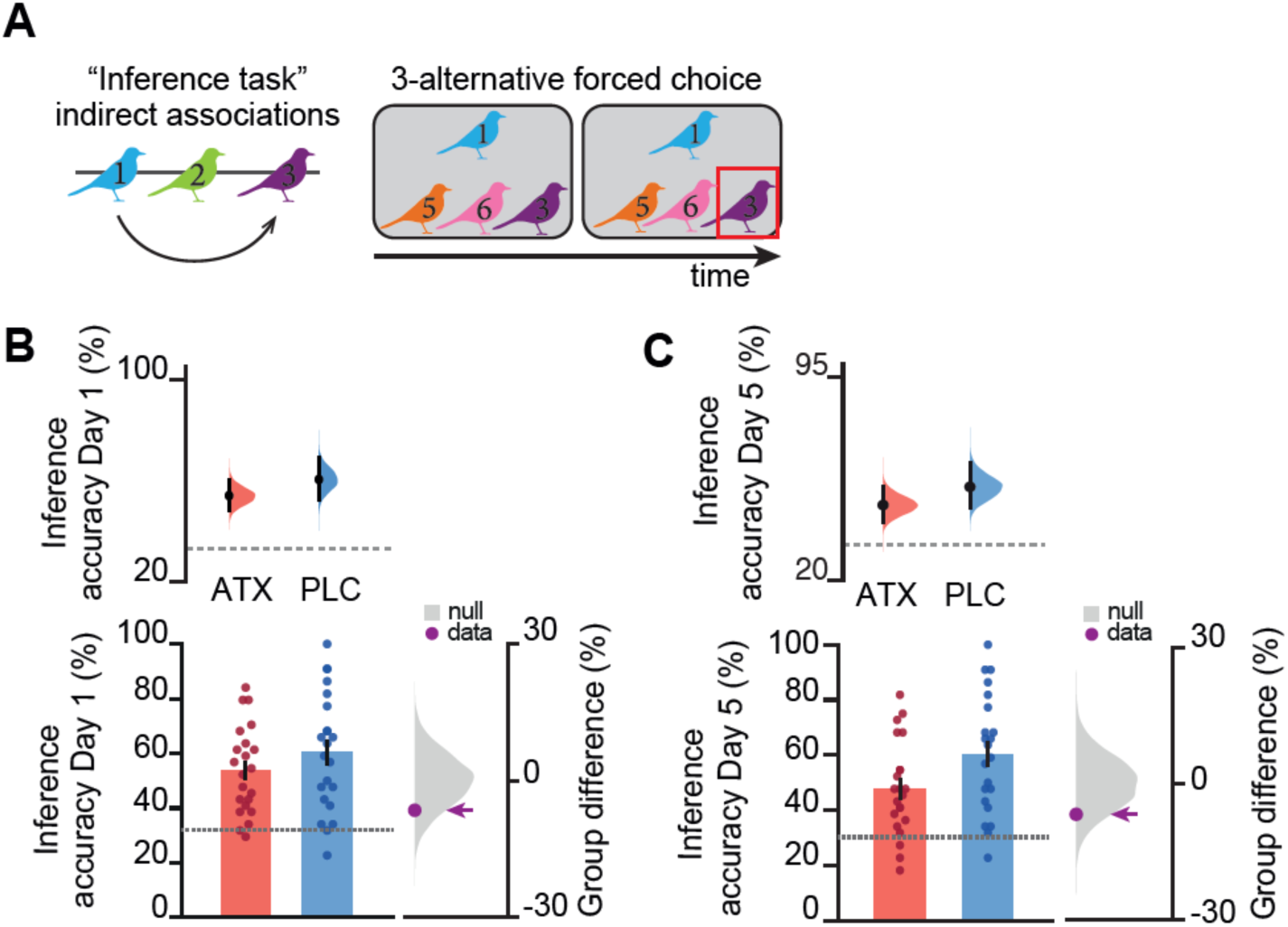
Inference task performed in day 1 and 5. **A** After the learning task (Figure 1D), participants inferred indirect relationships between bird stimuli in the absence of feedback, to encourage cohesive map learning (‘inference task’). **B** There was no significant difference in performance between groups on the inference task (ATX – PLC: permutation test: p=0.128). **C** On Day 5 participants repeated the inference task. Again, there was no significant difference in inference task performance between the groups (ATX – PLC: permutation test: p=0.124). **B-C** Upper: Bootstrap-coupled estimation (DABEST) plots. Black dot, mean; black ticks, 95% confidence interval; filled curve, sampling error distribution. Lower: memory accuracy (mean +/− SEM). Lower right: null distribution of the group differences generated by permuting subject labels, purple dot: true group difference.

**Supplementary Figure 4.**
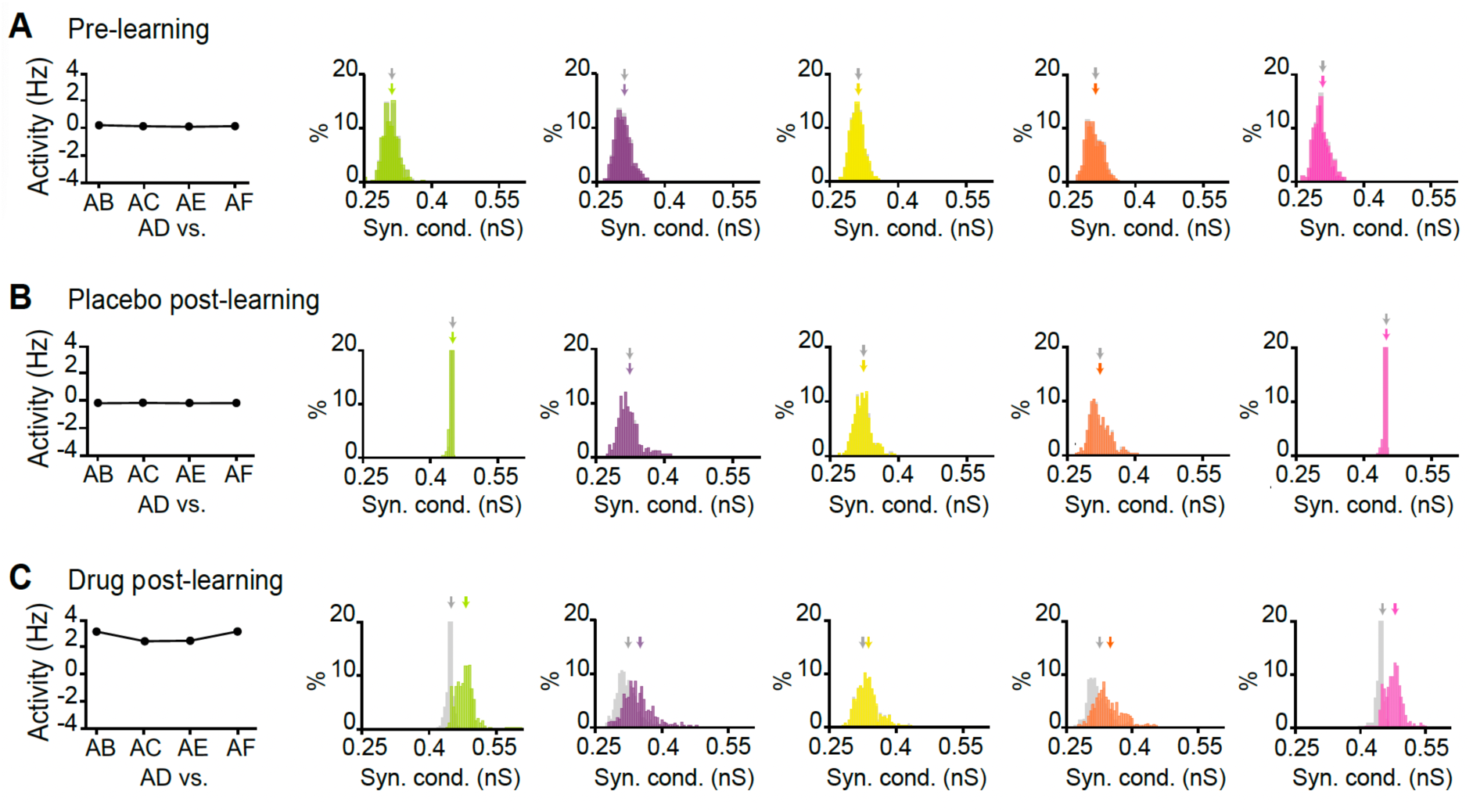
Additional features of the neural network model. Additional features of the neural network model under three different conditions: pre-learning (A), placebo post-learning (B) and atomoxetine post-learning (C). Shown are the average firing rate responses to stimulation of assembly ‘A’, together with the distribution of synaptic weights**. Left panel:** Proxy generated by the neural network for XSS contrast used to assess representational overlap with fMRI (Figure 7). Each plot shows the difference in average firing rate over the 3.3 s ‘stimulus’ period for assembly ‘A’ versus the 3.3 s stimulus period for another assembly (‘B’, ‘C’, E’ or ‘F’), relative to the average firing rate over the 3.3 s stimulus period for assembly ‘A’ versus assembly ‘D’ (the most distal assembly from ‘A’). For example, for ‘AB’ we show: [(average of ‘A’ – average of ‘D’) - (average of ‘A’ – average of ‘B’)]. **Rightward panels:** Distributions of weights of excitatory synapses between assembly ‘*A’* and all other assemblies (B: green; C: purple; D: yellow; E: orange; F: pink), 1 second before (grey) and 1 second after (coloured) the 3.3 s ‘stimulus’ period. Arrows indicate the mean value of each distribution. **A-B** In the pre-learning condition and in the post-learning placebo condition, the activities of all assemblies are uniform with respect to activity in *‘D’,* due to the lack of associations between assemblies (A) and sufficient inhibitory rebalancing (B). **C** Under atomoxetine, a gradient can be observed in the activities of assemblies with respect to activity in D, analogous to the XSS effect observed in fMRI data (Figure 7). **Rightward panels:** Distributions of weights of excitatory synapses between assembly ‘*A’* and all other assemblies (B: green; C: purple; D: yellow; E: orange; F: pink), 1 second before (grey) and 1 second after (coloured) the 3.3 s ‘stimulus’ period. The arrows indicate the mean value of each distribution.

**Supplementary Figure 5.**
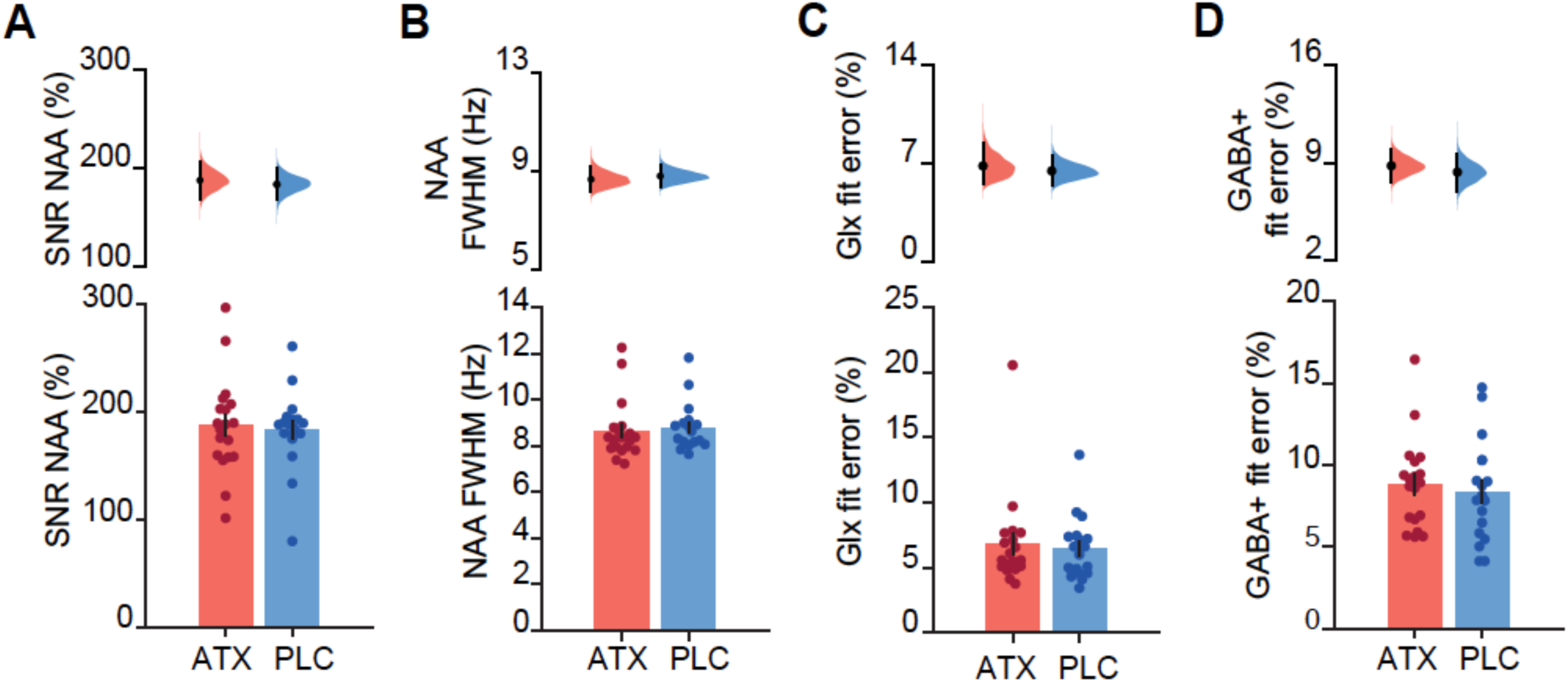
MRS quality measures for LOC showed no significant difference between groups. **A** The average signal-to-noise ratio (SNR) of N-acetylaspartate (NAA), a reference metabolite, as determined by Gannet (see Methods) was for 187.1% (SEM 10.24) for the ATX group and 183.2% (SEM 8.39) for the PLC group. There was no significant difference between groups (permutation test: p= 0.388). **B** The average full-width-at-half-max (FWHM) of NAA as determined by Gannet was 8.638 Hz (SEM 0.296) for ATX and 8.777 Hz (SEM 0.240) for PLC. There was no significant difference between groups (permutation test: p=0.363). **C** The average fit error for Glx as determined by Gannet was 6.820% (SEM 0.835) for ATX and 6.447% (SEM 0.563) for PLC. There was no significant difference between groups (permutation test: p=0.379). **D** The average fit error for GABA+ as determined by Gannet was 8.814% (SEM 0.632) for ATX and 8.344% (SEM 0.710) for PLC. There was no significant difference between groups (permutation test: p=0.312).

**Supplementary Figure 6.**
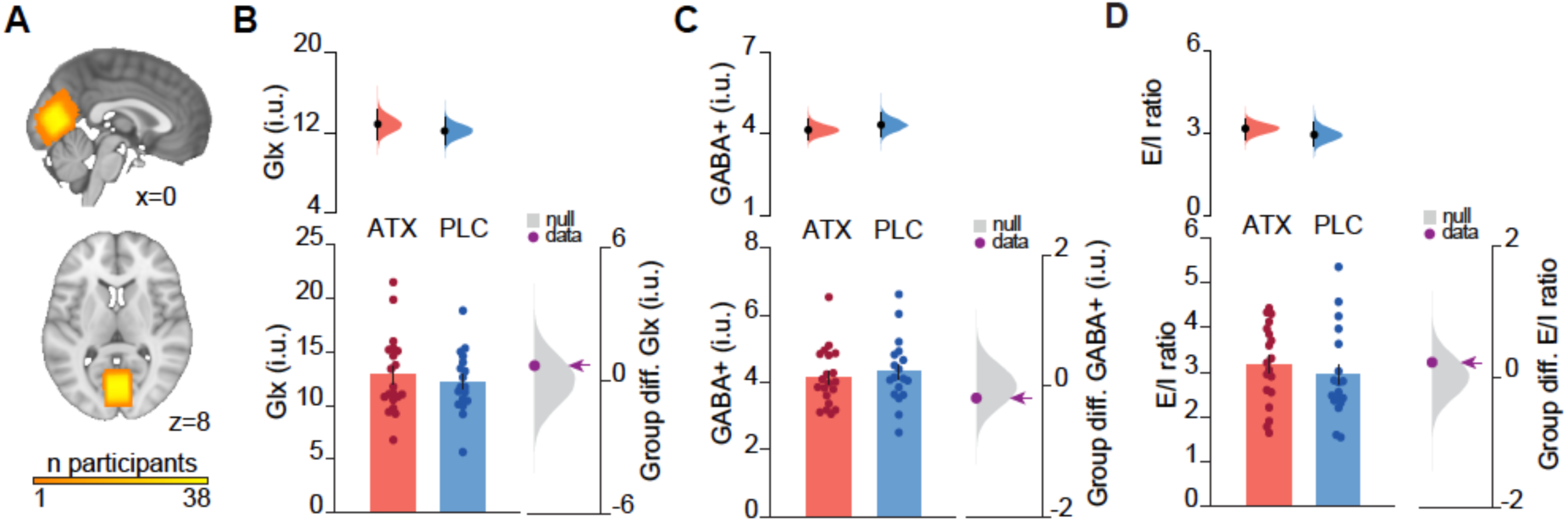
Elevating noradrenaline gives no significant change in Glx or GABA+ in V1. In contrast with the significant group difference reported in LOC (Figure 4), no significant difference in inhibitory processing was observed in V1. This may be explained by reduced SNR in V1 compared to LOC (Supplementary Figure 7), which may be attributed to spatial inhomogeneities in the static magnetic field (B0) or higher physiological noise levels in V1 (Hutton et al., 2011). **A** Anatomical location of the 2 x 2 x 2 cm^3^ MRS VOI, positioned in V1. Cumulative map across 38 participants. **B-D** We used MRS to quantify the effect of atomoxetine on excitatory and inhibitory tone in V1, ~3.5 h after drug intake. Upper: Bootstrap-coupled estimation (DABEST) plots. Black dot, mean; black ticks, 95% confidence interval; filled curve, sampling error distribution. Lower left: metabolite concentrations and E/I ratio (mean +/− SEM). Lower right: null distribution of the group differences generated by permuting subject labels, purple dot: true group difference. **B** No significant group difference was observed in the concentration of Glx (ATX – PLC: permutation test p=0.267). **C** No significant group difference was observed in the concentration of GABA+ (ATX – PLC: permutation test p= 0.269). **D** No significant group difference was observed in E/I ratio, defined as the ratio of Glx:GABA+ (ATX – PLC: permutation test p=0.233).

**Supplementary Figure 7.**
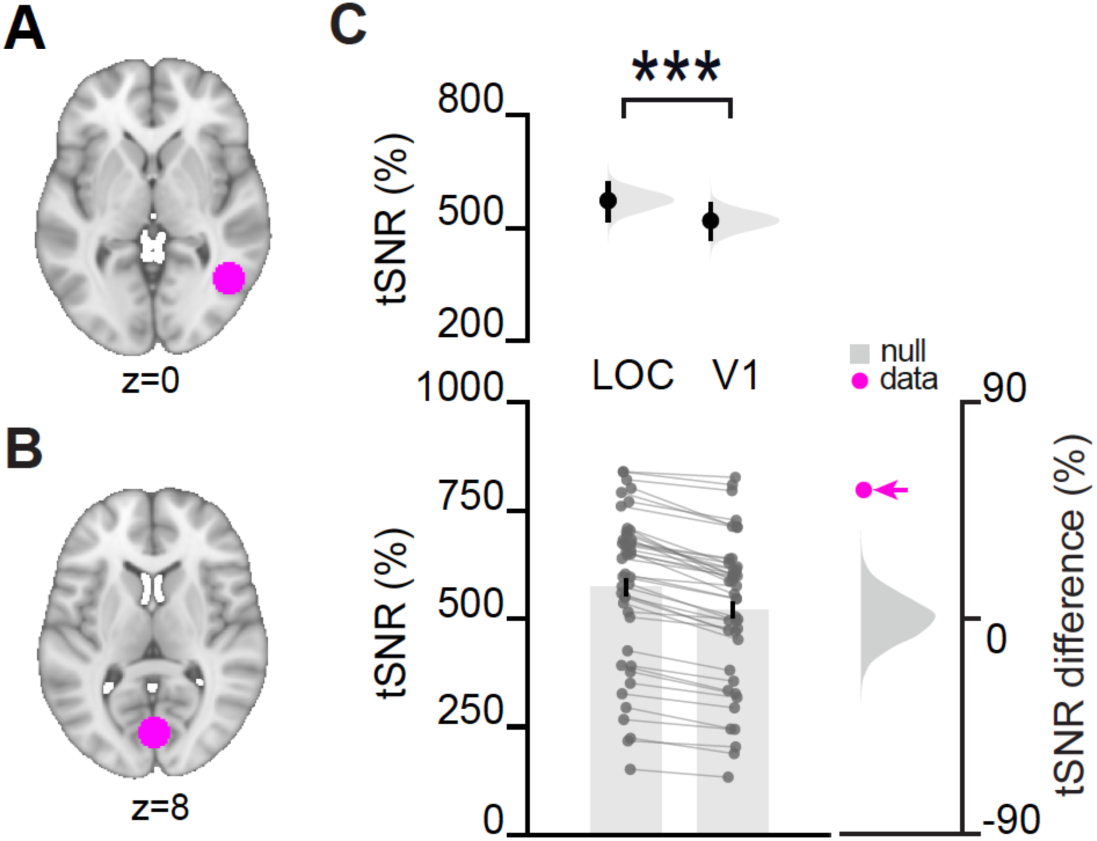
Higher signal to noise ratio (SNR) in LOC compared to V1. **A-B** ROIs used to determine tSNR and small volume correction. **A** 10mm radius sphere centered on the group-average MRS voxel positioned in LOC (Figure 4A). **B** 10mm radius sphere centered on the group-average MRS voxel positioned in V1 (Supplementary Figure 6A). **C** The temporal SNR (tSNR) calculated over the duration of the fMRI acquisition was significantly higher in LOC compared to V1 (permutation test: p<0.001). tSNR reflects MR signal quality and is influenced by factors that include the static magnetic field (B0) and physiological noise levels (Hutton et al., 2011). Upper: Bootstrap-coupled estimation (DABEST) plots. Black dot, mean; black ticks, 95% confidence interval; filled curve, sampling error distribution. Lower right: grey: null distribution of the tSNR differences; purple dot: true regional tSNR difference.

**Supplementary Figure 8.**
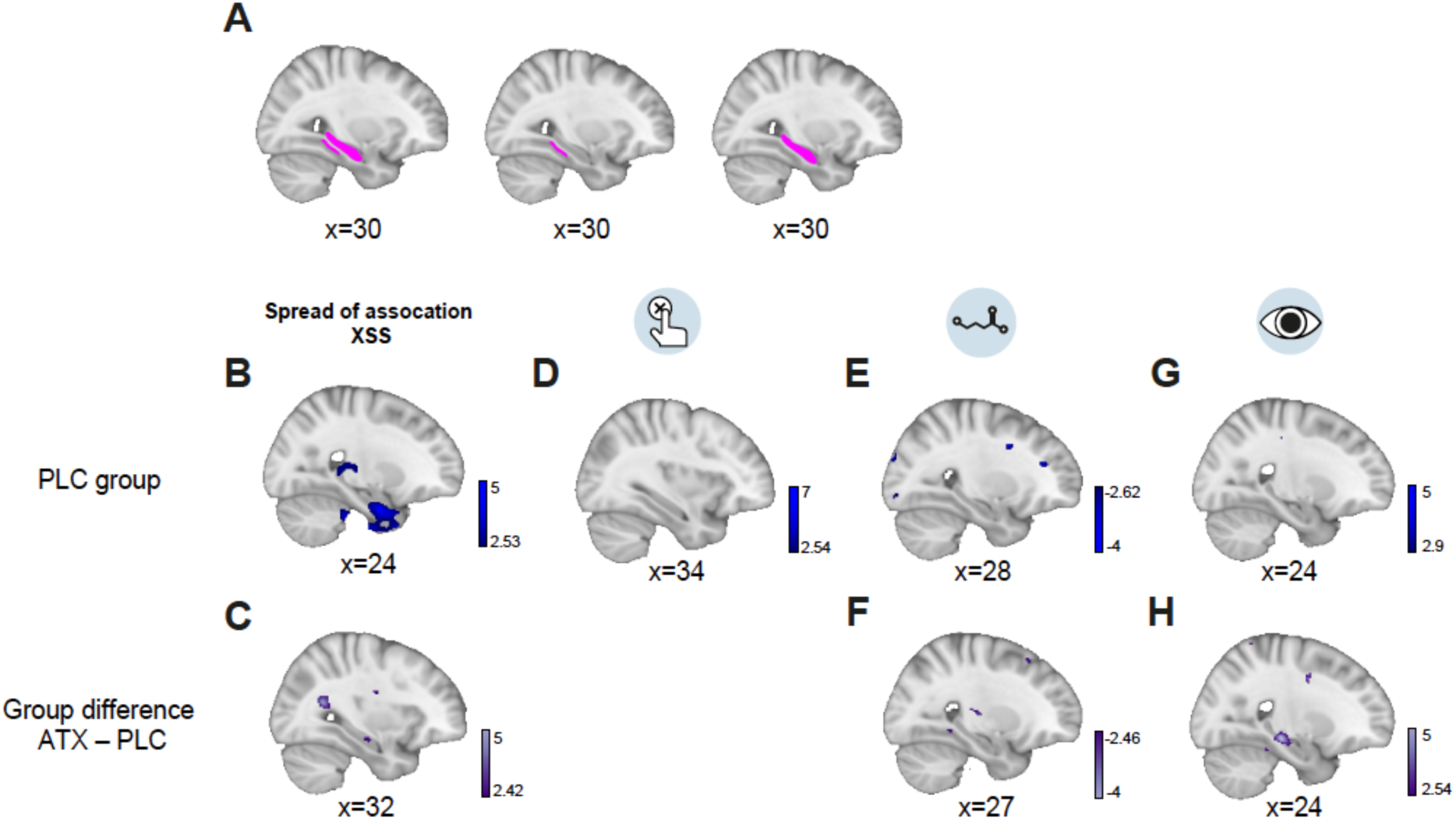
Regions of interest, placebo group effects and group differences for the neural spread of association results. **A** Left: Anatomical ROI of parahippocampus-hippocampus used for small volume correction. Middle and right: Anatomical ROIs of parahippocampus (middle) and hippocampus (right) used for *post hoc* tests. **B-H** In the PLC group, t-statistic maps showing the neural spread of association effect and correlations with this effect (B,D,E,G) and group differences between ATX and PLC (C,F,H). **B** In the PLC group, no significant effect was observed for the neural spread of association in the underlying memory map in parahippocampus-hippocampus, although a trend was observed (SVC with anatomical parahippocampus-hippocampus ROI, t_20_=3.85, p=0.122, MNI coordinates, Supplementary Table 3). **C** No significant group difference (ATX vs. PLC) was observed for the neural spread of association effect (SVC with parahippocampus-hippocampus ROI: t_41_=2.35, p=0.595, MNI coordinates). **D** In the PLC group, no significant relationship was observed between the neural spread of association and overgeneralisation errors reported in behaviour (SVC with parahippocampus-hippocampus ROI, t_19_=2.32, p=0.825, MNI coordinates). **E** In the PLC group, no significant correlation was observed between the concentration of GABA+ in LOC and the neural spread of association effect (SVC with parahippocampus-hippocampus ROI: t_14_=3.15, p=0.461). **F** No significant group difference (ATX vs. PLC) was observed for the relationship between GABA+ in LOC and the neural spread of association effect in hippocampus-parahippocampus (SVC with parahippocampus-hippocampus ROI: t_30_=3.17, p=0.195, MNI coordinates). **G** In the PLC group, no significant correlation was observed between the pupil dilation response to surprising stimuli and the neural spread of association effect (SVC with parahippocampus-hippocampus ROI: t_8_=3.63, p=0.501 MNI coordinates). **H** A trend towards a significant group difference (ATX vs. PLC) was observed for the relationship between the pupil dilation and the neural spread of association in parahippocampus-hippocampus (SVC with anatomical parahippocampus-hippocampus ROI: t_19_=3.45, p=0.078, MNI coordinates, Supplementary Table 8).

## Supplementary Tables

**Supplementary Table 1.** Effectiveness of blinding procedure Effectiveness of the blinding procedure assessed using Bang’s blinding index (BI) (see *Methods*, Bang et al., 2004). Importantly, the BI for ATX indicates successful blinding in this group, for both the participants and experimenter. The BI for the PLC of >0.2 for both participants and experimenter, indicating a tendency to correctly guess that PLC participants received placebo.

| Participants |  |  |  | Experimenter |  |  |  |
| --- | --- | --- | --- | --- | --- | --- | --- |
| Condition | BI | Confidence interval |  | Condition | BI | Confidence interval |  |
|  |  | Lower bound | Upper bound |  |  | Lower bound | Upper bound |
| ATX | -0.18 | -0.59 | 0.23 | ATX | 0.00 | -0.42 | 0.42 |
| PLC | 0.36 | -0.03 | 0.75 | PLC | 0.64 | 0.31 | 0.96 |

**Supplementary Table 2.** In the ATX group, cross-stimulus suppression as an index for spread of association; related to Figure 7. Brain regions showing significant cross-stimulus suppression for the parametric spread of association effect, as described in Figure 6B. No brain regions survived FWE whole-brain correction at the cluster-level (p<0.05). Statistics reported in the parahippocampus-hippocampus (Supplementary Figure 8A) are peak-level FWE corrected using a small-volume correction method. Post-hoc tests in parahippocampus and hippocampus (Supplementary Figure 8A) are peak-level FWE corrected using a small-volume correction method, with significant results reported (Figure 7A,B).

| Brain region | P<sub>FWE-corr</sub> | T<sub>peak level</sub> | Coordinate |  |  |
| --- | --- | --- | --- | --- | --- |
|  |  |  | x | y | z |
| Parahippocampus & hippocampus | P=0.032 | 4.56 | 22 | -36 | -18 |
| Parahippocampus & hippocampus | P=0.048 | 4.35 | 28 | -13 | -16 |
| Parahippocampus | P=0.011 | 4.56 | 22 | -36 | -18 |
| Hippocampus | P=0.033 | 4.35 | 28 | -13 | -16 |

**Supplementary Table 3.**
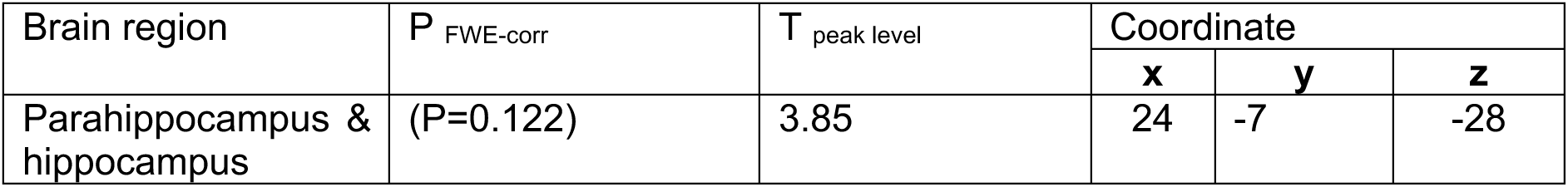
In PLC, cross-stimulus suppression as an index for the spread of association; related to Supplementary Figure 8. Brain regions showing cross-stimulus suppression for the parametric spread of association effect, as described in Figure 6B. No brain regions survived FWE whole-brain correction at the cluster-level (p<0.05). No brain regions survived FWE correction using a small-volume correction method, although a positive trend was observed in parahippocampus-hippocampus (Supplementary Figure 8A).

| Brain region | P<sub>FWE-corr</sub> | T<sub>peak level</sub> | Coordinate |  |  |
| --- | --- | --- | --- | --- | --- |
|  |  |  | x | y | z |
| Parahippocampus & hippocampus | (P=0.122) | 3.85 | 24 | -7 | -28 |

**Supplementary Table 4.** In the ATX group, correlation between cross-stimulus suppression as an index for spread of association and the behavioural measure of overgeneralisation errors; related to Figure 7. Brain regions showing significant correlation between cross-stimulus suppression for the spread of association effect (as described in Figure 6B) and the behavioural measure of overgeneralisation errors (as described in Figure 2F). No brain regions survived FWE whole-brain correction at the cluster-level (p<0.05). Statistics reported in the parahippocampus-hippocampus (Supplementary Figure 8A) are peak-level FWE corrected using a small-volume correction method. Post-hoc tests in hippocampus and parahippocampus are peak-level FWE corrected using a small-volume correction method, with significant results reported (Figure 7C).

| Brain region | P<sub>FWE-corr</sub> | T<sub>peak level</sub> | Coordinate |
| --- | --- | --- | --- |

|  |  |  | **x** | **y** | **z** |
| --- | --- | --- | --- | --- | --- |
| Parahippocampus & hippocampus | P=0.002 | 6.16 | 34 | -24 | -12 |
| Hippocampus | P=0.001 | 6.16 | 34 | -24 | -12 |

**Supplementary Table 5.** Group difference (ATX vs. PLC) for the correlation between cross-stimulus suppression as an index for spread of association and the behavioural measure of overgeneralisation errors; related to Figure 7. Brain regions showing a significant group difference (ATX – PLC) in correlation between cross-stimulus suppression as an index for the spread of association (as described in Figure 6B) and overgeneralisation errors reported in behaviour. No brain regions survived FWE whole-brain correction at the cluster-level (p<0.05). Statistics reported in the parahippocampus-hippocampus are peak-level FWE corrected using a small-volume correction method. Post-hoc tests in hippocampus and parahippocampus (Supplementary Figure 8A) are peak-level FWE corrected using a small-volume correction method, with significant results reported (Figure 7D).

| Brain region | P<sub>FWE-corr</sub> | T<sub>peak level</sub> | Coordinate |  |  |
| --- | --- | --- | --- | --- | --- |
|  |  |  | **x** | **y** | **z** |
| Parahippocampus & hippocampus | P=0.011 | 4.51 | 36 | -22 | -14 |
| Hippocampus | P=0.007 | 4.51 | 36 | -22 | -14 |

**Supplementary Table 6.**
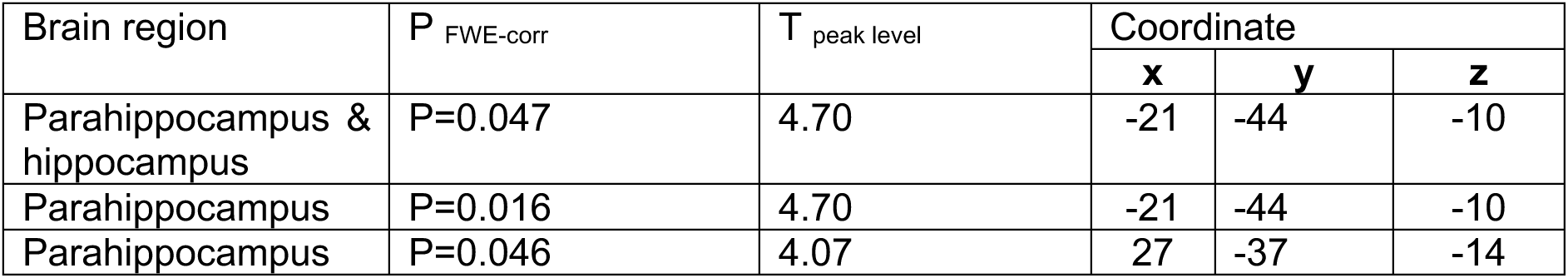
In the ATX group, correlation between GABA+ in LOC and cross-stimulus suppression as an index for spread of association; related to Figure 8. Brain regions showing significant correlation with GABA+ in LOC and cross-stimulus suppression as an index for spread of association in the underlying memory map (as described in Figure 6B). No brain regions survived FWE whole-brain correction at the cluster-level (p<0.05). Statistics reported in parahippocampus-hippocampus (Supplementary Figure 8A) are peak-level FWE corrected using a small-volume correction method. Post-hoc tests in hippocampus and parahippocampus (Supplementary Figure 8A) are peak-level FWE corrected using a small-volume correction method, with significant results reported (Figure 8A).

| Brain region | P<sub>FWE-corr</sub> | T<sub>peak level</sub> | Coordinate |  |  |
| --- | --- | --- | --- | --- | --- |
|  |  |  | **x** | **y** | **z** |
| Parahippocampus & hippocampus | P=0.047 | 4.70 | -21 | -44 | -10 |
| Parahippocampus | P=0.016 | 4.70 | -21 | -44 | -10 |
| Parahippocampus | P=0.046 | 4.07 | 27 | -37 | -14 |

**Supplementary Table 7.** In the ATX group, correlation between pupil dilation response to surprising stimuli and cross-stimulus suppression as an index for spread of association; related to Figure 8. Brain regions showing significant correlation between the pupil dilation response to surprising stimuli and cross-stimulus suppression as an index for the spread of association effect (as described in Figure 6B). No brain regions survived FWE whole-brain correction at the cluster-level (p<0.05). Statistics reported in the parahippocampus-hippocampus (Supplementary Figure 8A) are peak-level FWE corrected using a small-volume correction method. Post-hoc tests in hippocampus and parahippocampus (Supplementary Figure 8A) are peak-level FWE corrected using a small-volume correction method, with significant results reported (Figure 8B).

| Brain region | P<sub>FWE-corr</sub> | T<sub>peak level</sub> | Coordinate |  |  |
| --- | --- | --- | --- | --- | --- |
|  |  |  | **x** | **y** | **z** |
| Parahippocampus & hippocampus | (P=0.066) | 5.15 | 21 | -18 | -16 |
| Hippocampus | P=0.045 | 5.15 | 21 | -18 | -16 |

**Supplementary Table 8.** Group difference (ATX vs. PLC) for the correlation between cross-stimulus suppression as an index for spread of association and the pupil dilation response to surprising stimuli; related to Supplementary Figure 7. Brain regions showing a trend group difference (ATX – PLC) in correlation between the pupil dilation response to surprising stimuli and cross-stimulus suppression as an index for the spread of association effect (as described in Figure 6B). No brain regions survived FWE whole-brain correction at the cluster-level (p<0.05). Statistics reported in the parahippocampus-hippocampus (Supplementary Figure 8A) are peak-level FWE corrected using a small-volume correction method.

| Brain region | $P_{\text{FWE-corr}}$ | $T_{\text{peak level}}$ | Coordinate | | |
| --- | --- | --- | --- | --- | --- |
|  |  |  | **x** | **y** | **z** |
| Parahippocampus & hippocampus | ($P=0.078$) | 3.45 | -27 | -24 | -19 |

